# Architecture of Near-Death Experience Spaces

**DOI:** 10.1101/2025.10.17.682526

**Authors:** France Lerner, Guillaume Tahar, Netta Shafir

## Abstract

Near-Death Experiences (NDEs) challenge conventional research methods, as they are often described as exceeding the limits of verbal expression. While most studies rely on narrative accounts, little attention has been given to graphic representations as a complementary mode of investigation. This study employs a hybrid approach combining a semi-structured digital questionnaire with a graphic reconstruction task. Participants first segmented their Out-of-Body Experience (OBE) and NDE into distinct experiential sequences by assigning chronological numbers to each phase. This sequencing enabled the identification and isolation of the core NDE component across participants. The graphic reconstructions represent each perceived space, Self-Location (S-L), the perceived spatial position where one experiences oneself to be located and Self-Motion (S-M), the perceived spatial trajectories and directionalities of oneself within these visuo-spatial configurations. This dual-method design was intended to capture experiential features of OBEs and NDEs that may not be conveyed through verbal reports alone. Notably, OBEs have rarely been examined regarding their independence from or conjunction with NDEs, nor their placement within the overall temporal sequence. The graphic reconstructions revealed consistent visual-field invariants: conical forms, elliptic arcs with variable Visual-Field Extents (VFE), and ellipsoidal configurations, each correlated with specific chromatic and luminance perceptual qualities of the Visual Field (VF).

## 1 Introduction

NDEs are profound, often transformative experiences that typically occur in life-threatening situations such as cardiac arrest (Parnia et al. 2001), respiratory failure, or coma. Phenomenologically similar experiences have also been reported in non-lethal contexts (Owens et al. 1990; Charland-Verville et al. 2014), including brain disorders (Facco and Agrillo 2012b), post-traumatic coma (Hou et al. 2013), states of emotional shock (Greyson 2014), recreational drug use (e.g., N,N-Dimethyltryptamine, DMT), and altered states of consciousness such as meditation or REM sleep (Peinkhofer et al. 2021). In such cases, these are commonly referred to as NDE-like. Both types of NDEs are frequently characterized by vivid multisensory mental imagery and are often perceived as subjectively real. (Thonnard et al. 2013; Palmieri et al. 2014; Martial et al. 2019), prompting interdisciplinary interest across neuroscience, psychology, and phenomenology to develop explicative models (Fritz et al. 2024).

Recent studies estimate that 4–8% of the general population has had an NDE, with a frequency up to 10% in a cross-cultural sample of 1,034 individuals across 35 countries (Kondziella et al. 2019). Incidence is higher among clinical populations, with 6–23% of cardiac-arrest survivors reporting such experiences (Van Lommel et al. 2001; Schwaninger et al. 2002; Klemenc-Ketis 2013). Although most reports arise in life-threatening contexts, an estimated 15–30% occur in non-life-threatening contexts (Charland-Verville et al. 2014). Despite their prevalence, NDEs remain difficult to investigate due to ineffability often linked to emotional intensity (Thonnard et al. 2013). Although NDE memories resemble real memories phenomenologically and electrophysiologically, exhibiting temporal theta activity consistent with episodic recall (Moore and Greyson 2017), their vivid perceptual and affective content may exceed the expressive capacity of verbal reports, supporting the use of alternative methodologies to access and represent spatial and visual structure.

Among commonly reported features, OBEs hold a distinctive place. OBEs are characterized by a sense of disembodiment in which an illusory body, distinct from the physical one, is perceived, and the sense of Self-Location (S-L) is altered such that the visuospatial perspective is taken from a parasomatic position. As Blanke (2012) defines it, “self-location, that is, the experience of where I am in space” is typically at the location of the body and closely tied to the first-person perspective; in OBEs this S-L is transposed to an extracorporeal space. Neurophysiological evidence from focal electrical stimulation of the right angular gyrus suggests such experiences may result from a breakdown in the brain’s integration of complex somatosensory and vestibular signals, leading to altered whole-body perception and displacement (Blanke and Arzy 2005; Blanke et al. 2002).

In the general population, 10–20% report at least one OBE (Blackmore 1984; Murray and Fox 2005). In the context of NDEs, 45–80% report OBEs during the episode (Greyson 2014; Van Lommel et al. 2001; Parnia et al. 2014), typically early in the NDE and often coinciding with observing one’s physical body from an elevated, external viewpoint. These frequencies suggest OBEs may constitute a recurrent component of NDEs phenomenological structure. Although phenomenological studies have identified recurring features such as tunnel perception, encounters with deceased individuals, and out-of-body perspectives (Zaleski 1987; Blackmore 1993; Belanti et al. 2008; Greyson et al. 2009; Greyson 2015), less attention has been directed to the visuo-spatial structure of these experiences and to how individuals visually perceive and kinesthetically experience their Self-Motion (S-M) within them.

The study of NDEs progressed from early philosophical and theological inquiry to empirical investigation within the human and medical sciences. The first systematic collection was published by St. Gallen Heim (1892), and the term “near-death experience” was introduced Egger (1896) in *The Self of the Dying*. Early explorations by parapsychologists, including Sir William Barrett *(Death-Bed Visions, 1926)*, were met with skepticism, delaying integration into rigorous academic research. Interest reemerged in the late 20th century through foundational contributions in psychology and psychiatry by (Moody, 1975; Ring, 1980; Greyson, 1984; Sabom, 1982).

With advancements in neuroscience, explanatory models have been developed in neurology (Vanhaudenhuyse et al. 2009; Blanke et al. 2016), neurochemistry (Martial et al. 2019), neurophysiology (Morse et al. 1989), and neurobiology (Saavedra-Aguilar and Gómez-Jeria 1989); Britton and Bootzin (2004). Electrophysiological studies have investigated correlates of NDE-like experiences using EEG during unconscious or altered states: (Martial et al. 2019) examined the neurophenomenology of NDE memories via hypnotic recall, and (Martial et al. 2024) identified EEG signatures during syncope-induced unresponsiveness. These studies suggest altered connectivity among visual cortex, temporo-parietal junction, and default mode network.

Historically, both Western and non-Western artistic depictions have represented altered states, including NDEs (León 2013). Iconographic parallels appear in Bosch’s *Visions of the Hereafter* (*Ascent of the Blessed*, 1500–1504), Fuseli’s *The Nightmare* (1781), and Doré’s *Vision of the Empyrean* (1868). Modern cinema reiterates NDE motifs in *The Wizard of Oz* (Fleming, 1939), *Flatliners* (Schumacher, 1990), and *Brainstorm* (Trumbull, 1983). For OBEs, both Western artistic traditions (e.g., Schiavonetti after William Blake’s *The Soul Hovering Over the Body*, 1805–08) and non-Western forms such as Aboriginal Australian painting (e.g., Western Desert works by Clifford Possum Tjapaltjarri) depict elevated, extracorporeal viewpoints.

Prior investigations emphasized written reports, thematic categorization, and artistic representations (Kennedy 1997; Rominger 2004; Tang 2019), without addressing the sequential organization of OBE spaces in relation to NDE spaces, the chronological progression of NDE spatial configurations, or the evolving dynamics of S-L and S-M.

Our study builds upon previous research, including earlier investigations into the spatial organization of OBEs and NDEs (Lerner 2016; Lerner et al. 2021), by introducing structured sequencing of visuo-spatial configurations and systematically mapping S-L, S-M, and VMI, we aim to identify recurring VF invariants in OBEs and NDEs. We interpret the VFs represented in NDErs graphic representations as instances of what (Haun and Tononi 2025) describe as the, intrinsic structure of visual experience: “It is not necessarily the representation of the space of the outside world (commonly referred to as visual space). The spatial structure of the visual field is largely invariant to the organization of the world it mediates.”

In our study, each NDEr’s boundary outline graphical representation represents the intrinsic spatial structure of the VF as experienced during the NDE. The VF is understood as the spatial form of visual awareness during the experience. Further, the graphic representation are mapping NDERs first-person perspective self spatial dynamics within the VF boundary. We analyzed the marked S-L and S-M relative to each VFE outline to determine how these factors influenced the curvature of the VF boundary and its VFE. Ultimately, we aim to establish a systematic framework for analyzing the spatial organization of OBEs and NDEs, contributing to a more precise characterization of their experiential structure and content.

## 2 Materials and methods

### 2.1 Ethical approval

This study was approved by the Ethics Committee of the Unit for Interdisciplinary Studies, Bar-Ilan University, Tel Aviv (IRB No. ISU202208001). All participants provided informed consent, and the study adhered to ethical standards protecting participants’ rights and privacy.

### 2.2 Participants

Eligibility required fluency in English, age *≥* 18, and absence of mental disability or guardianship. Classification as a Near-Death Experiencer (NDEr) followed completion of a preliminary questionnaire including two validated instruments: the Greyson NDE Scale (Greyson 1983) and the NDE Content Scale (NDE-C; (Martial et al. 2020)) (ESM, Preliminary Questionnaire). The Greyson Scale (16 items, 0–2; total 0–32) defines NDEs with scores *≥* 7. The NDE-C (20 items, 0–4; total 0–80) confirms NDErs with scores *≥* 27.

Among 104 volunteers (69 female, 33 male; *M* = 54.7, *SD* = 14), those meeting both score cut-offs (*≥* 7 Greyson; *≥* 27 NDE-C) were invited to the second phase of the study to answer the main questionnaire. Participants completed the study online on a voluntary basis, with the option to withdraw at any time. All data were anonymized, and drawings were converted to grayscale using Adobe Photoshop 2024 (version 25.4).

### 2.3 Methodology

Our study characterized NDEs and OBEs perceptual structures as experienced (Husserl 1982), this approach emphasizes the perspectival, horizon-structured, and embodied nature of visual experience (Gibson 2014; Thompson 2010; Madary 2016). First-person reports were analyzed to identify recurrent NDEs VF types and perceptual features. These categories were then operationalized as quantitative variables, following principles of experimental phenomenology by merging qualitative descriptions with systematic quantification of perceptual properties (Albertazzi et al. 2011; Albertazzi 2013). This integration parallels experimental neurophenomenology, in which first-person accounts are treated as structured data suitable for coding and statistical analysis (Thompson, 2010), thus ensuring reproducibility and comparability.

### 2.4 Questionnaire design and administration

A semi-structured, researcher-assisted digital questionnaire was administered remotely via Google Meet. It consisted mainly of closed-ended, checkbox-style items ensuring response consistency and quantitative analysis, supplemented by open-ended questions to capture qualitative details (ESM, Main questionnaire 1–4; 5–7; 8–10). Before the interview, participants received an instructional email outlining the procedure and were asked to print a reference sheet defining key terms and symbols to be used during the session.

The questionnaire, implemented in Google Forms, was completed concurrently with the graphic tasks. The reference document contained the catalog of NDE pictograms, the lexicon and the graphic symbols used in the graphic reconstruction tasks (ESM, Catalog), along with examples of OBE Dissociative and Associative phases and NDE space types A–C (ESM, Examples). During the live session, the researcher guided participants and provided clarifications as needed, the session duration ranged from 1.5 to 3 hours, depending on the experiential complexity of the participant’s OBE and/or NDE.

### 2.5 Instructions for recalling and structuring OBE and NDE

Participants were encouraged to take as much time as needed to recall their OBE and/or NDE and were invited to close their eyes if this facilitated memory retrieval. The researcher then reviewed the lexicon with each participant to ensure full understanding of all terms. After this clarification phase, participants were asked to divide their experience into sequential parts, which could include an OBE, an NDE, or both. To aid segmentation, metaphorical analogies were offered such as scenes in a film, acts in a play, chapters in a book, or movements in music and participants selected whichever best suited their narrative style.

Each participant first identified whether the experience contained an OBE component. When present, the OBE was classified as *Dissociative*, *Associative*, or both. A Dissociative OBE involved adopting a parasomatic viewpoint and perceiving the physical body and/or the surrounding physical space while experiencing a felt separation from it. In contrast, an Associative OBE involved the same parasomatic viewpoint and perception of the physical body and/or physical space, but was accompanied by a felt re-embodiment into it. Each OBE type counted as one part; if both occurred, they were recorded separately, yielding two OBE parts. Participants then ordered their graphic reconstructions chronologically, the Dissociative OBE preceding the NDE sequence and the Associative OBE following it. The number of printed pages equalled the number of parts, summarized by the formula: “X parts = Y OBE parts + Z NDE parts.”

Finally, participants numbered each part in chronological order and reviewed the full sequence, including OBE components and NDE Core parts, before advancing to the next stage.

### 2.6 Graphic reconstruction task

For the graphic representation tasks of OBEs and NDEs, participants used a blue pen (1.00 mm tip). For the OBE reconstructions, participants first reviewed illustrative examples (ESM, *Examples*, pp. 1, 5), then drew a single outline to represent the physical environment in which the OBE occurred and annotated it with standardized symbols: a circle for the physical body, a cross for S-L, an arrow for the direction of S-M, and squares for other beings (e.g., medical staff or animals).

For the NDE reconstructions, participants first reviewed illustrative examples (ESM, *Examples*, pp. 2–4) then selected from the catalog the predefined NDE pictogram that best matched the perceived boundary of the space they saw or experienced themselves as being within during their NDE. (ESM, *Catalog*, pp. 1–2). The chosen pictogram was redrawn with precision to reflect individual perception. If the same space was experienced more than once, the same pictogram could be reused; if none matched, participants produced a freehand drawing. Using standardized graphical symbols, participants annotated drawings to indicate S-L and S-M for each sequence. S-L were marked with numbered crosses (*×*), e.g., S-L1 and S-L2, indicating their chronological order; S-M was represented with arrows (*→*) showing directional trajectories, with U-shaped arrows indicating reversals in direction. Squares marked the position of other entities, physical beings in OBEs and non-physical presences (e.g., spiritual guides, angels, or deceased relatives) in NDEs (ESM, *Catalog*, p. 4). All drawings followed a fixed spatial convention: the bottom of the page represented the ground or physical floor, and the top indicated the ceiling, sky, or upward extension, maintaining a vertical frame aligned with gravitational orientation.

### 2.7 Digital questionnaire data collection

The digital questionnaire was implemented in Google Forms and completed in parallel with the graphic reconstruction tasks. For each sequence depicted in their drawing, participants kept the graphic representation in view while answering the corresponding questionnaire items. This ensured that the visual record they had just produced directly informed their categorical and descriptive responses, reducing recall bias and supporting accurate mapping between drawn and verbal content.

The questionnaire comprised both closed-ended check-box items and open-ended text fields. The open-ended section elicited rich qualitative descriptions of each NDE space sequence. To preserve the authenticity of these descriptions, only minor spelling corrections were made. The closed-ended section systematically recorded, for each spatial sequence: (i) the numbers of S-L, (ii) the S-M velocity profiles, (iii) the emotional valence and intensity ratings for each S-L and (iv) the sensory/perceptual modalities experienced in the NDE space (e.g., visual, auditory, haptic, etc.) The graphic representations described the NDE spatial boundary, the spatial positioning and sequential numbering of the S-L, the directional trajectory of the S-M, and the spatial locus of the mental imagery (e.g., other beings). In the present study, only variables (i) and (ii) were analyzed, and within the verbal descriptions we examined only the perceptual appearance of the NDE spaces specifically their chromaticity and luminance qualities rather than the full narrative content.

#### Supplementary data and reference conventions

All supporting text-based and visual materials used in this study are provided as Electronic Supplementary Material (ESM). These include the catalog of NDE pictograms, participant graphic representations, illustrative examples of OBE and NDE spatial configurations, questionnaires (preliminary and main), and NDE sequences, showing how participants divided and chronologically ordered the components of their experience (with OBE phases when present). In the main text and discussion, references to these materials use the abbreviation “ESM” followed by the specific document name (e.g., ESM, graphic representation). Supplementary Material appearing at the end of the manuscript is referred to in full as “Supplementary Material A, B or C,” to distinguish it from the ESM documents.

## 3 Results

The results are presented in four sections. (1) We distinguished NDEs from NDE-like experiences based on participants’ self-attributed etiology. (2) We examined the distribution of OBE components (Dissociative, Associative, or both) and the categorization of NDE spatial forms into A-, B-, and C-shapes. (3) We analyzed the geometric characteristics of NDE VFs, focusing on VFE. (4) We analyzed the relationship between S-L, S-M, and VFE outlines to determine how these factors influenced the perceived curvature of the VF boundary.

### 3.1 Distinction of NDE and NDE-like experiences based on self-attributed etiology

In this analysis, subject numbers denote anonymized identifiers. The dataset includes only the 50 participants who completed the main questionnaire, excluding those who participated exclusively in the preliminary one. Participants were asked whether they considered their NDE to have occurred under life-threatening conditions. Based on self-reports, 42 participants (84%) judged their experience as life-threatening and were classified as NDErs, while 8 (16%) described it as non-life-threatening, corresponding to the NDE-like category. This distribution indicates that most NDEs in this subset were perceived as occurring under life-threatening circumstances.

### 3.2 Distribution of OBE and NDE shape categories (A, B, C)

Fifty subjects completed the main digital questionnaire and produced graphic representations according to its protocol. They were asked to specify the number of parts in their NDE, indicate whether they experienced an OBE, and locate it chronologically within the NDE. Participants were also instructed to distinguish between a Dissociative OBE (subjective impression of separating from the physical body) and an Associative OBE (subjective impression of reuniting with the physical body). Among the 50 participants, 18 reported only a NDE without any OBE component (Subjects 2, 3, 16, 18, 22, 23, 25, 26, 27, 28, 31, 32, 35, 36, 38, 43, 45, 48). Eleven described an OBE Dissociative component with an NDE but not the Associative type (Subjects 1, 4, 7, 8, 13, 17, 19, 37, 41, 47, 50). Nine reported an OBE Associative component but not the Dissociative type (Subjects 9, 10, 12, 15, 20, 21, 40, 46, 49). Twelve experienced both Dissociative and Associative OBEs within the same sequence (Subjects 5, 6, 11, 14, 24, 29, 30, 33, 34, 39, 42, 44). This distribution shows that OBEs are frequent but not universal within NDEs, with Dissociative and Associative types occurring in nearly equal measure, and some participants experiencing both as transitional phases within the broader NDE structure (Supplementary Material A, Figure 4).

Table shows the distribution of the number of NDE sequence reported by the 50 subjects. The median number of segments is 2, while the mean is 2.24.

**Table 1:** Distribution of the number of NDE parts per subject (*n* = 50). The median number of segments is 2, and the mean is 2.24.

| Number of parts | 1 | 2 | 3 | 4 | 5 | 6 | 7 | 8 |
| --- | --- | --- | --- | --- | --- | --- | --- | --- |
| Count | 21 | 14 | 7 | 3 | 2 | 2 | 0 | 1 |
| Relative frequency | 42% | 28% | 14% | 6% | 4% | 4% | 0% | 2% |

For the 21 participants who reported a single NDE sequence, the distribution of the selected shapes is presented in Table 2 (Supplementary Material), while all possible shape options are shown in the Catalog (ESM, pp. 1–2).

**Table 2:** Distribution of shapes for subjects with a single NDE part (*n* = 21).

|  | A | B | C1–4 | C5 | Other (freehand) | No space |
| --- | --- | --- | --- | --- | --- | --- |
| Count | 3 | 3 | 8 | 5 | 1 | 1 |
| Relative frequency | 14.3% | 14.3% | 38.1% | 23.8% | 4.8% | 4.8% |

For the OBE, Table 3 (Supplementary Material A) presents the S-M orientations for the 10 participants who experienced both an Associative and a Dissociative OBE with rectilinear trajectories. The angles were measured in a reference frame where 0° corresponds to the horizontal axis pointing right and 90° to the vertical axis pointing upward on the page, and the last column reports the difference between these two angles (invariant under a change of reference frame).

**Table 3:** S-M orientations (rectilinear) in Dissociative vs. Associative OBEs. Angles measured in a 2D page-fixed frame: 0*^◦^*= rightward horizontal, 90*^◦^* = upward vertical. Subject 24, whose S-Ms were rotational, is not included in this table even though they reported both an Associative and a Dissociative OBE.

| Subject ID | Dissociative orientation ($^\circ$) | Associative orientation ($^\circ$) | Relative angle ($^\circ$) |
| --- | --- | --- | --- |
| 5 | 90.0 | -90.0 | -180.0 |
| 11 | 45.7 | -90.0 | -135.7 |
| 14 | 90.0 | -90.0 | -180.0 |
| 29 | 56.0 | -39.7 | -95.7 |
| 30 | 62.7 | -149.5 | -212.2 |
| 33 | 21.2 | 178.0 | 156.8 |
| 34 | 90.0 | -90.0 | -180.0 |
| 39 | 122.2 | -40.6 | -162.8 |
| 42 | 79.4 | -99.0 | -178.4 |
| 44 | -131.7 | -166.3 | -34.6 |

We suggest that A shapes tend to appear at the beginning or end of the series. To test this, we examined NDEs with more than one part and measured how often each shape type appeared in the middle (i.e., not first or last). The proportion of middle occurrences among multi-part series was significantly lower for A shapes than for the other groups. Table 4 (Supplementary Material A) shows, for each shape type (A, B, C1–4, and C5), the proportion of middle appearances and the corresponding t-statistics and p-values comparing each group to A shapes. Results are significant at *p <* 0.016 after Bonferroni correction.

**Table 4:** Relative frequency of shapes occurring in middle positions of NDE sequences (only sequences with *>* 1 part). Lower panel: one-sample *t*-tests comparing each ratio to the A-shape ratio. Bonferroni-corrected significance threshold *p <* 0.016.

|  | A | B | C1-4 | C5 |
| --- | --- | --- | --- | --- |
| Total occurrences (seq. $> 1$ part) | 25 | 28 | 12 | 24 |
| Middle occurrences | 1 | 16 | 7 | 9 |
| Relative frequency | 4% | 57.1% | 58.3% | 37.5% |
| $t$-statistic vs. A | — | -5.145 | -3.53 | -3.085 |
| $p$-value | — | 0.002 | 0.002 | 0.000005 |

For the C5 shapes, 18 participants reported more than one shape and their shape sequences included C5. Table 4 (Supplementary Material) shows which shapes appeared before and after C5, and Table 5 (Supplementary Material A) provides the corresponding frequencies.

**Table 5:**
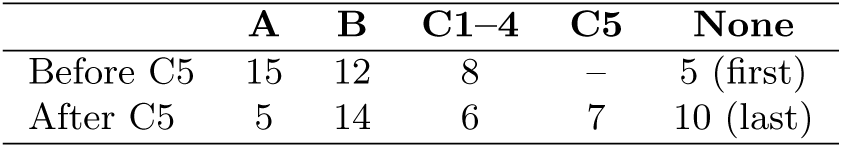
Distribution of shapes preceding and following C5. “None” indicates C5 at sequence boundary (first/last).

|  | A | B | C1–4 | C5 | None |
| --- | --- | --- | --- | --- | --- |
| Before C5 | 15 | 12 | 8 | – | 5 (first) |
| After C5 | 5 | 14 | 6 | 7 | 10 (last) |

To further examine the relationships between different NDE space types and OBE components, we analyzed their co-occurrence in the same experiential episodes, without considering their sequential order (Supplementary Material A, Figure 4)

### 3.3 NDE VF geometric characteristics

Based on the classification of space types (Supplementary Material B), we quantified the geometric properties of VF boundaries relative to S-L. In each drawing, participants marked their S-L within the depicted space. For A, B, and C1–4 shapes, the VFE was defined as the angle between the S-L (or the first, when multiple) and the two edges of the shape. For A shapes, we additionally defined an opening angle the angle between the two boundary lines of the shape.

Figure 1 illustrates these definitions: the right panel shows the opening angle *θ* between the two boundary lines of an A shape, representing the enclosed VFE; the left panel shows an ellipsoidal shape with a S-L, where *θ* is the angular extent between the S-L and the two edges, indicating the perceived range of vision. These measures enabled quantitative comparison of visual perspectives reported by NDErs across spatial configurations.

**Fig. 1:**
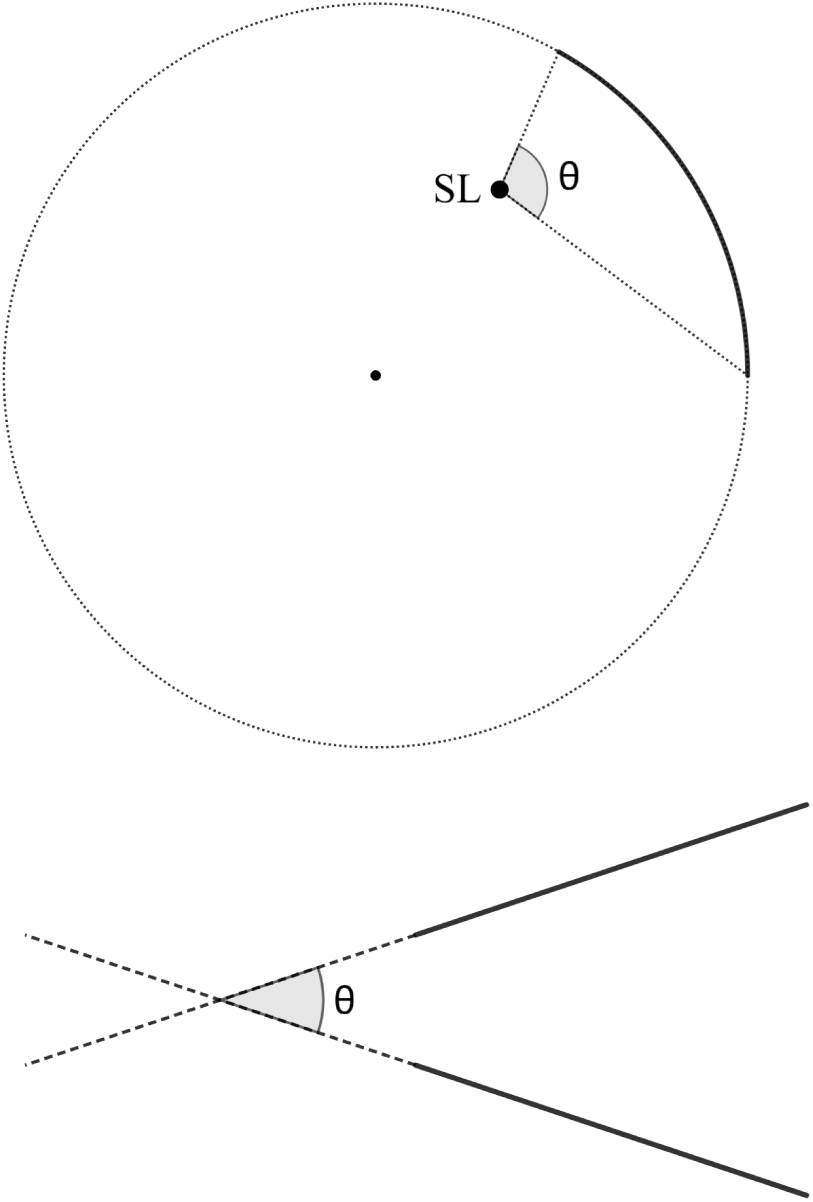
Measurement of Visual Field Extent (VFE) and opening angle in NDE graphic representations. right: opening angle *θ* for an A shape; left: ellipsoidal shape with S-L (SL), where *θ* spans the two edges visible from SL.

**Fig. 2:**
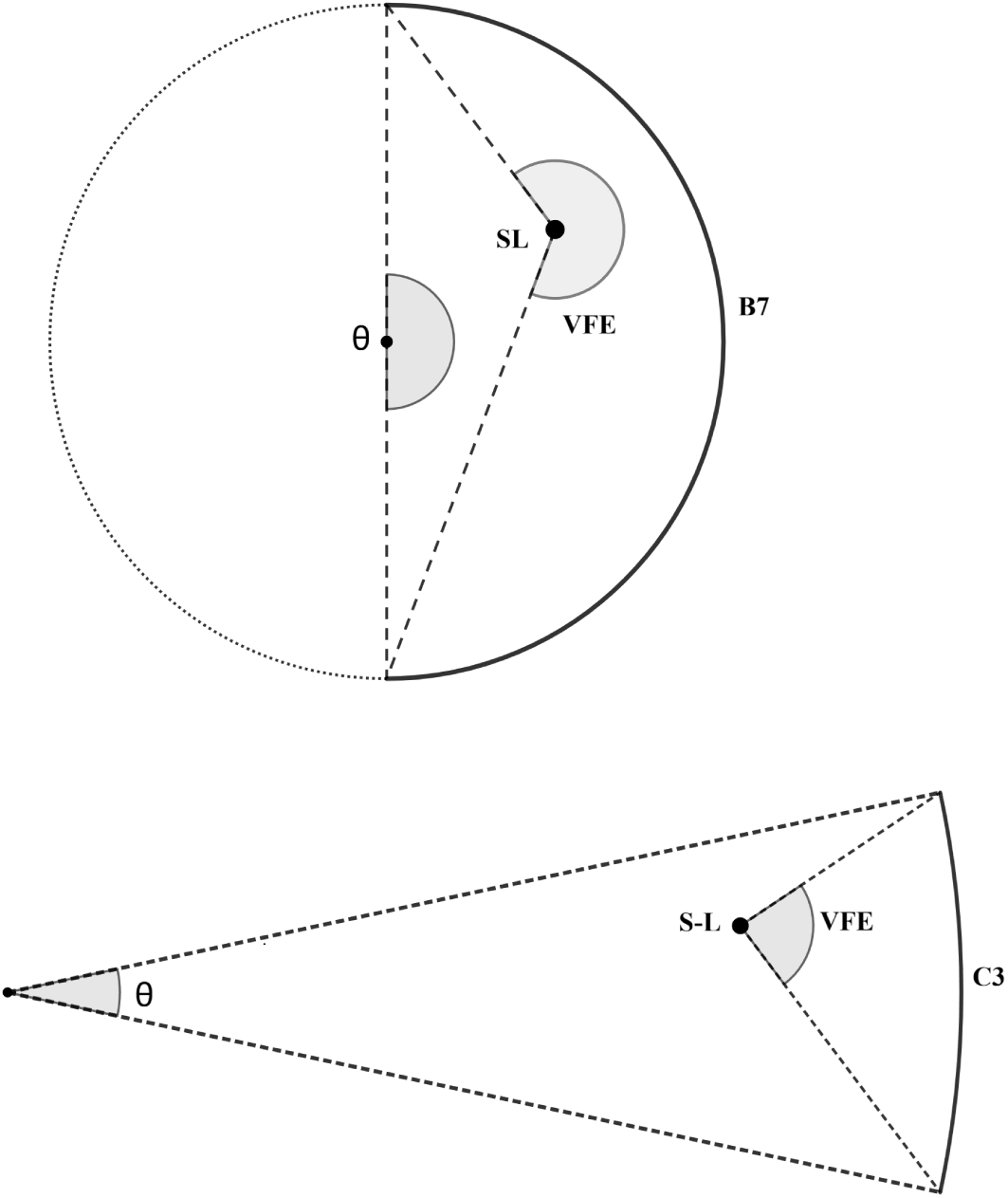
Examples illustrating how curvature and S-L distance modulate VFE. Up: B7 with high angular amplitude and pronounced curvature; down: C3 with low curvature and reduced angular amplitude.

**Fig. 3:**
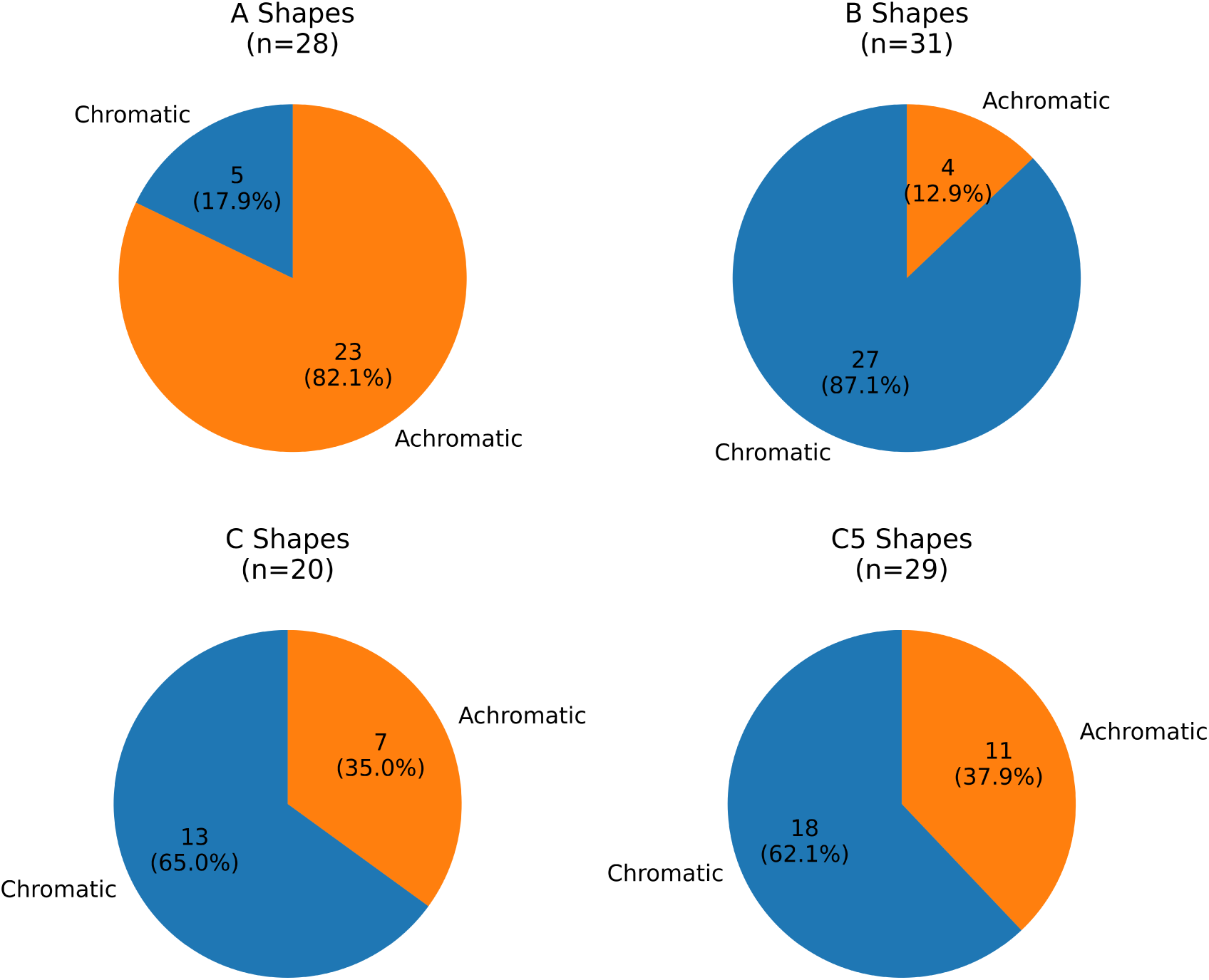
Distribution of color palette (chromatic vs. achromatic) by shape category (A, B, C1–4, C5).

For A shapes, the distribution of opening angles is shown in Figure 5 (Supplementary Material A). Figure 6 (Supplementary Material A) presents the distributions of VFEs for B shapes and C1–4 shapes (excluding C5). In both figure panels, box plots display the median, and histograms display the mean (orange lines).

**Fig. 4:**
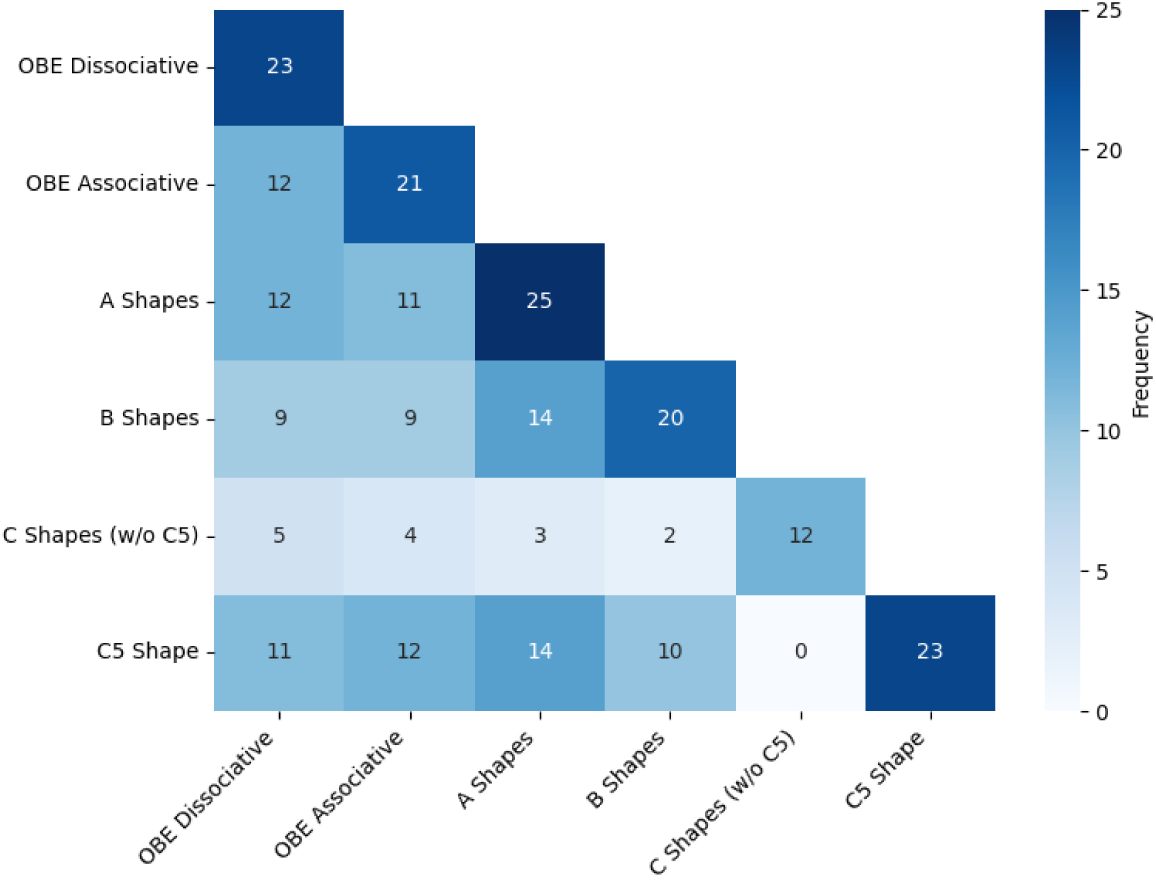
Co-occurrence matrix of NDE shape categories (A1–12, B1–12, C1–4, C5) and OBE parts.

**Fig. 5:**
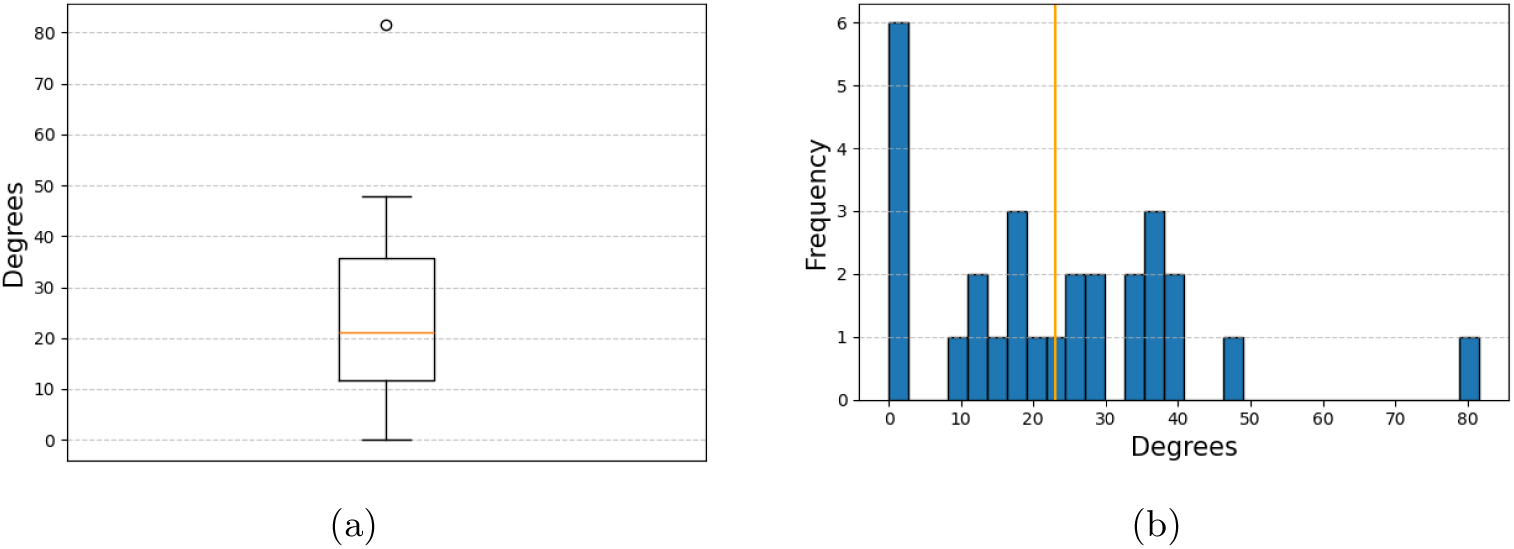
Distribution of opening angles for A shapes. (a) Box plot showing the median; (b) histogram showing the mean.

**Fig. 6:**
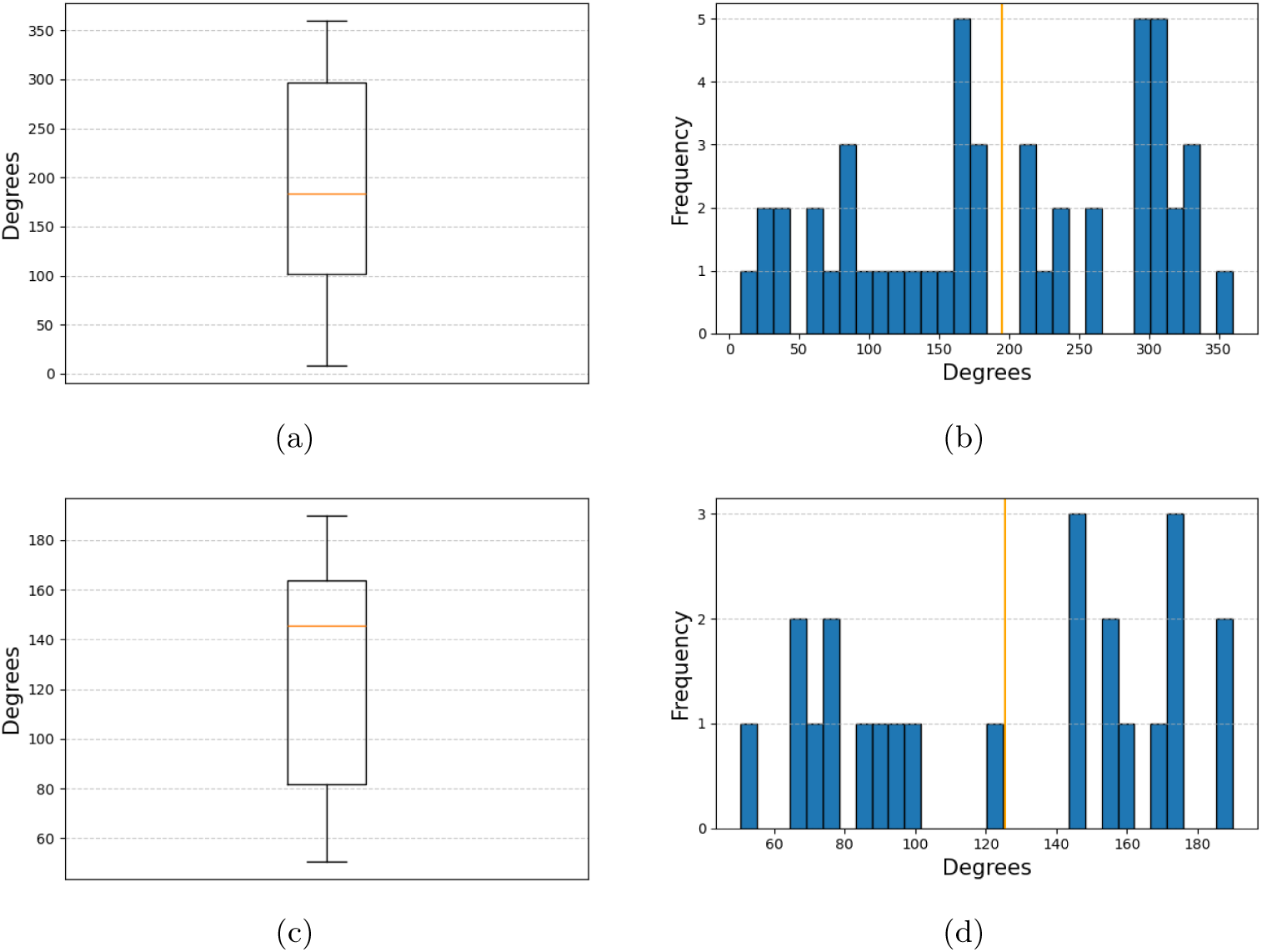
Distribution of VFEs for B and C1–4 shapes. (a) Box plot of VFEs in B shapes; (b) histogram of VFEs in B shapes; (c) box plot of VFEs in C1–4 shapes; (d) histogram of VFEs in C1–4 shapes.

We additionally examined whether the opening angle of an A shape correlated with the VFE of the S-L in a subsequent ellipsoidal shape (B or C1–4) within the same NDE sequence. Across all such paired occurrences, the correlation was *−*0.133, indicating no strong linear relationship between these two measures.

Figure 2 left panel illustrates B7, where the arc has a high angular amplitude and pronounced curvature. The S-L lies relatively close to the arc, resulting in a larger VFE. In contrast, the right panel illustrates C3, where the arc curvature has low-curvature and the angular amplitude is reduced, approaching a flat profile. Here, the S-L is located farther from the arc relative to its radius of curvature, yielding a smaller VFE. Together, these examples demonstrate how both curvature and S-L distance modulate the orientation and extent of the VFE.

### 3.4 NDE VF perceptual qualities and MI

Figure 3 presents the distribution of chromatic versus achromatic appearances across shape categories, with C5 treated separately. A-shapes (*n* = 28) were mostly achromatic (23 cases, 82.1%), whereas B-shapes (*n* = 31) were predominantly chromatic (27 cases, 87.1%). C-shapes (*n* = 20) showed a more balanced pattern (13 chromatic, 65%; 7 achromatic, 35%), and C5 spaces (*n* = 29) were primarily chromatic (18 cases, 62.1%). For C5, the achromatic group included Subjects 1, 5, 10, 14, 17, 20, 21, 25, 42, 43, and 48; the chromatic group included Subjects 6, 11, 18, 22, 23, 39, 40, 41, 43, 44, 45, 46, 47, and 21.

## 4 Discussion

Most studies on NDEs have not systematically examined how the experience unfolds by characterizing specific VF types. In contrast, the present study uses participants’ segmentation of their NDEs into discrete sequences, each linked to a distinct visuo-spatial configuration. This enables a chronologically ordered analysis of invariant VF features and perceptual transitions across the temporal course of the experience.

Recent large-scale citizen-science projects most notably the Perception Census led by Anil Seth and colleagues (Perception Census Team 2022) have validated freehand participant drawing as a scientific tool for studying subjective visual phenomenology under controlled stimulation. As part of the Dreamachine project, more than 15,000 visual representations were collected to document individual differences in perception elicited by identical white-light stroboscopic sequences. Building on this methodological precedent, the present discussion develops a multidimensional analysis of the spatio-phenomenological structure of Out-of-Body Experiences (OBEs) and NDEs, integrating participants’ graphic reconstructions with their written reports through a descriptive phenomenological approach.

The discussion is organized around four key points emerging from the analysis:

1. Sequencing of OBE and NDE experiences;
2. Differentiation between OBE and NDE;
3. NDE VF boundaries;
4. Chromatic qualities within the VF.

### 4.1 Sequencing OBEs and NDEs: isolating the structure of the experiential NDE Core

OBEs commonly occur within NDEs but may also appear independently, implying distinct causal pathways. Conversely, NDE-like experiences phenomenologically similar to canonical NDEs (e.g., tunnel vision, light, life review) can arise in non-life-threatening contexts such as emotional shock, syncope, or psychedelic use (Fritz et al. 2024). This challenges the view that NDEs depend solely on proximity to death, suggesting shared neurocognitive mechanisms, including REM dysregulation and altered temporoparietal activity. Among the 50 participants completing the main questionnaire, 42 (84%) reported life-threatening conditions (classified as NDErs), and 8 (16%) described NDE-like experiences in non-life-threatening contexts. The sequencing task enabled participants to divide their experience into temporally ordered phases, delineating a NDE Core.

After reviewing the lexicon (ESM, *Catalog*, p. 3), participants segmented their experience assigning to each phase a sequential label with its corresponding pictogram (ESM, *Catalog*, pp. 1–2). OBEs were consistently structured into two components Dissociative and Associative visually encoded using standardized symbols: a circle for the physical body, a cross for S-L, arrows for S-M direction, and squares for entities. The Dissociative OBE was characterized by upward or lateral S-M away from the body, whereas the Associative OBE involved a downward S-M with a sense of re-embodiment. A list of subjects and corresponding graphic representations for each phase is provided in (Supplementary Material A, Table 6).

**Table 6:** Subjects and pages for Dissociative vs. Associative OBE phases (from Supplementary Material: Graphic Representations).

| Subject ID | Page | OBE Phase |
| --- | --- | --- |
| 1 | 3 | Dissociative |
| 4 | 14 | Dissociative |
| 5 | 18 | Dissociative |
| 5 | 21 | Associative |
| 6 | 23 | Dissociative |
| 6 | 30 | Associative |
| 7 | 32 | Dissociative |
| 8 | 36 | Dissociative |
| 9 | 41 | Associative |
| 10 | 48 | Associative |
| 11 | 50 | Dissociative |
| 11 | 59 | Associative |
| 12 | 65 | Associative |
| 13 | 67 | Dissociative |
| 14 | 71 | Dissociative |
| 14 | 74 | Associative |
| 15 | 78 | Associative |
| 17 | 82 | Dissociative |
| 19 | 91 | Dissociative |
| 20 | 98 | Associative |
| 21 | 106 | Associative |
| 24 | 115 | Dissociative |
| 24 | 117 | Associative |
| 29 | 132 | Dissociative |
| 29 | 136 | Associative |
| 30 | 138 | Dissociative |
| 30 | 140 | Associative |
| 33 | 147 | Dissociative |
| 33 | 149 | Associative |
| 34 | 151 | Dissociative |
| 34 | 153 | Associative |
| 37 | 158 | Dissociative |
| 39 | 164 | Dissociative |
| 39 | 166 | Associative |
| 40 | 169 | Associative |
| 41 | 171 | Dissociative |
| 42 | 175 | Dissociative |
| 42 | 179 | Associative |
| 44 | 184 | Dissociative |
| 44 | 186 | Associative |
| 46 | 192 | Associative |
| 47 | 194 | Dissociative |
| 49 | 203 | Associative |
| 50 | 205 | Dissociative |

Participants’ graphic representations revealed consistent differences in S-M orientation between the two OBE phases. S-M was measured using a shared reference frame, with 0*^◦^* defined as horizontal S-M to the right of the page. The relative angle between Dissociative and Associative orientations was independent of this convention. Across subjects, relative angles were typically close to 180*^◦^*, reflecting opposite orientations between the two OBEs (Supplementary Material A, Table 3). Exceptions, such as Subject 24, involved rotational S-Ms of 90*^◦^*, geometrically conjugated by 180*^◦^* rotation (ESM, *Graphic Representations*, pp. 115–117).

The sequencing protocol instructed participants to position Dissociative OBEs at the onset and Associative OBEs at the conclusion of their NDE sequencing. Participants endorsed this ordering as consistent with their lived temporal dynamics, enabling consistent isolation of a NDE Core, the experiential sequence(s) situated between the two OBEs. The resulting tripartite framework: Dissociative OBE *→* NDE Core *→* Associative OBE provided a coherent structure for cross-subject spatial analyses (ESM, *Sequences*). The overall co-occurrence between Dissociative and Associative OBEs and the NDE shape categories A1–12, B1–12, C1– 4, and C5 is summarized in (Supplementary Material A, Figure 6), which visualizes their frequency of joint appearance across all experiential sequences.

### 4.2 The differentiation of OBEs and NDEs

A key empirical distinction between OBEs and NDEs in our study lies in how participants represented the physical body and its environment. In OBE reconstructions particularly during the Dissociative phase both body and spatial context (e.g., room or location) were systematically depicted, showing that bodily visibility and spatial grounding were integral to the experience. These depictions indicate that the self was perceived as extracorporeally located yet still visually tethered to the body and its surroundings (Blanke 2004; Blanke et al. 2016) In contrast, during NDEs participants consistently refrained from depicting either the body or the surrounding physical environment, despite explicit instruction that such representation was possible. This absence suggests a phenomenological decoupling in which body and environment cease to serve as reference points for S-L. The distinction can be clarified through the framework of Haun and Tononi (2019), who state: “The term space can be taken to refer to a variety of concepts: there is the physical space of the world that we take to exist independent of our knowledge of it (‘outer space’); there is spatial cognition, the psychological model of physical space that an animal builds and uses to navigate and understand the world; and there is phenomenal space, the fact that some aspects of experience have a spatial structure.” Interpreted in this context, participants in OBEs retain a representation of physical space with a reference frame anchored by the body, whereas in NDEs this anchoring disappears, revealing a shift toward phenomenal space alone. This empirical contrast, the presence versus absence of bodily representation, the basis for isolating a NDE Core set of invariant features.

In NDEs, the physical body is often reported as no longer felt but replaced by a self lived as a viewpoint in a first-person perspective. The OBE–NDE transition can thus be seen as a shift of embodiment from the physical body now perceived as insentient to a self experienced as a viewpoint in a first-person perspective, marking a transfer of corporeal awareness from somatic to visuo-vestibular systems, consistent with Lucci and Pazzaglia (2015).

To illustrate this distinction, two representative graphic reconstructions were contrasted. The first is a Dissociative OBE followed by an NDE space of any ABC type; in one example, the Dissociative OBE is followed by a conical A1 shape (ESM, Graphic Representations, Subject 5, p. 18 for the OBE Dissociative and p. 19 for A1). The second example shows an Associative OBE preceded by a conical A5 shape (ESM, Graphic Representations, Subject 9, p. 40 for A5 and p. 41 for the OBE Associative). Together, these examples highlight a progressive shift in self-perception: in OBEs, the experiential viewpoint transitions from the physical body (represented by a circle) to a parasomatic S-L (represented by a cross). In NDEs, the parasomatic viewpoint becomes the sole referent for self-perception. S-L is no longer anchored to the physical body but is consistently experienced at the first-person perspective across all ABC-type sequences, defining the NDE Core and its associated visuo-spatial invariants.

### 4.3 Invariant geometries of Core NDE spaces VF

The categorization into the A-shape family was based on geometric properties of participants’ selections, with all A-shapes conceived as conical forms differing primarily in orientation either parallel or convergent (ESM, Catalog p. 1). The geometric parameters used to describe these VF configurations are defined in Figure 1, which illustrates the measurement of the opening angle (*θ*) in conical A-shapes and the VFE in ellipsoidal configurations. A key finding is that while the geometry of these spaces remains invariant, changes in S-M direction or S-L rotation can make the same shape appear inverted or rotated. For instance, in Subject 10, the sequence progressed from A5 to A9 through spaces B6, C5, and B7. A9 appears as the image of A5 under a 180° rotation (ESM, Sequences p. 5; Graphic Representations, A5 p. 42; A9 p. 46). This inversion reflects the reversal of S-M direction, while the underlying conical geometry remains constant. Another example, from Subject 23, shows a transition from A9 to A8 through C5 (ESM, Sequences p. 8). Here, the A-shape rotates rather than inverts, reflecting a rotation of S-L within C5. The graphic representation shows A9 (ESM, Graphic Representations p. 111) becoming A8 under a 225° clockwise rotation (ESM, Graphic Representations p. 113), with a residual trace of A9’s outline visible along the edge of C5 (ESM, Graphic Representations p. 112). Together, these cases demonstrate that the conical geometry of A-shapes is preserved as an invariant feature of Core NDE spaces, even when inversions or orientation changes arise relative to preceding spatial frames. The distribution of these A-, B-, C1–4-, and C5-type geometries across subjects with single-part NDEs is summarized in (Supplementary Material A, Table 2).

The categorization into B and C shape families was likewise based on geometric properties of participants’ pictogram selections, all conceived as elliptic arcs except C5, which represents a complete ellipse. By construction, the angular amplitude and curvature are high for B1–12 and low (almost flat) for C1–4. The VFE increases as the distance between S-L and the arc decreases relative to the radius of curvature. Figure 2 illustrates how curvature and S-L distance jointly modulate the amplitude of the VFE.

The B1–12 and C1–4 forms correspond to smaller or larger arcs of the complete ellipse represented by C5. Participants tended to select B when perceiving themselves far from the VF boundary and C1–4 when close to it. For example, in Subject 20’s graphic representation, the S-L in B6 lies farther from the VF boundary than in the following C2 arc, where it is closer (ESM, Graphic Representations pp. 95–96). This suggests that B and C shapes can be conceived as partial arcs of the C5 ellipse, with S-L distance influencing perceived VFE size and curvature. The relative frequency of these shapes in middle positions of NDE sequences is reported in (Supplementary Material A, Table 4. Rotational shifts of S-L may reorient VF boundaries without altering intrinsic geometry (ESM, Graphic Representations, Subject 25, B12–C5–B10, pp. 119–121). Translational S-M may also modify perceived VFE structure. Subject 47’s spaces B10 and B11 (pp. 197–198) show a transition between two elliptic arcs where the VFE changes along a translational trajectory.

Empirically, C5 graphic representations exhibit varying ellipse eccentricities from nearly spherical or circular to distinctly ellipsoidal depending on the ratio of major and minor axes. More elongated ellipsoidal geometries appear in Subjects 1, 39, and 40 (ESM, Graphic Representations, p. 6; p. 165; p. 168), whereas nearly spherical forms are found in Subjects 5, 6, and 41 (ESM, Graphic Representations, p. 20; p. 29; p. 172). If B1–12 and C1–4 elliptic arcs derive from the complete C5 ellipsoid, their geometry may also vary with perceived C5 eccentricity, as illustrated in Subjects 6 and 12 (ESM, Graphic Representations, pp. 25–26; pp. 63–64). These rare exceptions confirm the robustness of the pictogram methodology for representing NDE VF boundaries (ESM, Catalog pp. 1–2).

Participants’ chosen pictograms closely matched their verbal descriptions of perceived VF boundaries. Across cases, a consistent phenomenological invariant emerged: the VF assumed conical (A1–12), partial elliptic (B1–12; C1–4), or full ellipsoidal (C5) forms. These recurring geometries delineate the perceptual architecture of NDEs and support their systematic organization by VF boundaries and transformations distinguishing conical from partial or complete ellipsoidal structures (Figure 1). Such transformations serve as temporal anchors, segmenting the experience into perceptually defined phases. Consequently, NDE sequencing reflects transitions between successive VF boundaries, enabling temporal sequencing grounded in visuo-spatial configuration.

This framework differentiates OBE and NDE phenomenology, identifies invariant VF boundaries, and characterizes their chromaticity. It can also be extended to analyze additional experiential dimensions, including self-motion velocity profiles, sense of agency, and affective components. Sequential segmentation thus provides a robust basis for correlating experiential events with their visuo-spatial architecture.

### 4.4 Chromatic perceptual qualities across core NDE VFs

To examine the perceptual qualities of the VFs, we analyzed the distribution of chromatic (colorful) and achromatic (grayscale) features across all NDE Core spatial categories: A (A1–A12), B (B1–B12), C (C1–C4), and C5, treated here as a distinct category. The following discussion addresses the A category first, then B and C, and concludes with the C5 visuo-spatial configuration. Having established chromatic–achromatic distributions across A, B, C, and C5 categories (Figure 3), a central finding is the systematic variation of chromaticity across shape types. A1–12 shapes were predominantly achromatic (82.1%), suggesting reduced color perception within conical VF boundaries. In contrast, B1–12 shapes were largely chromatic (87.1%), indicating strong color perception within elliptic-arc VFs. C1–4 shapes showed a more balanced yet chromatic-dominant profile (65%), while C5 spaces displayed mixed proportions (62.1% chromatic, 37.9% achromatic).

#### Category of conical VF boundaries (A1–A12)

In our sample, 23 of 28 subjects (82.1%) experienced A-shaped spaces with predominantly achromatic perceptual qualities, while 5 (17.9%) reported chromatic ones. Participants’ accounts emphasized conical geometry and dark achromatic tones. For instance, Subject 5 (A1): “all the colors around me were rather dark, gray, black of different shades” (Supplementary Material C); Subject 6 (A1): “the space was silent and black …” (p. 3); Subject 7 (A1): “I was contained in a tubelike space …” (p. 4); Subject 13 (A5): “felt I was going through star fields; black. Saw stars that were bright but they were stationary; I was moving” (p. 7); Subject 15 (A5): “it looks like night time …” (p. 15); Subject 21 (A9): “escalator in a pure black space. Floating upwards through pitch blackness. 45 degrees, constant motion” (p. 9). Light modulation was salient in Subject 22 (A9): “moved through tunnel, started to lighten as I progressed, started black as night, finished as gray early dawn” (p. 10).

All 28 subjects reported achromatic perception except five (Subjects 4, 7, 10, 19, 42) where the A-shape appeared chromatic. In four, it occurred first in sequence; in Subject 42, second (ESM, Sequences, p. 12). These rare chromatic cases likely reflect perceptual overlap between consecutive spaces: when an A-shape preceded a B-shape, residual chromaticity from the latter may have been perceived while still within the A-shape. Two examples illustrate this (ESM, Sequences, Subjects 4 p. 4 and 10 p. 5).

For Subject 4, the sequence began with A1 (p. 15) followed by B10 (p. 16) (ESM, Graphic Representations, A1 p. 15; B10 p. 16). In A1, S-L faced the “garden of heaven”; the next space, B10, corresponded to the “pictures around wall of life.” On p. 16, B10 reappears with identical geometry. The participant described A1: “Dark to start … saw images on the walls of my life (life review) in vivid colour with sound, like a movie.” B10: “The being at the end was pure light … The garden was huge with rolling hills, tall grass … children playing, animals roaming.” This suggests that S-L within A1 sometimes allowed perceptual access to the next chromatic B10 panorama hosting life-review imagery. Similarly, Subject 10’s sequence began with A5 followed by B6, both showing transitional chromaticity. In A5 (p. 42), S-L lies at the end of the conical form; in B6 (p. 43), entities appear within the expanded field (ESM, Graphic Representations, A5 p. 42; B6 p. 43). A5: “moved very fast. Walls seemed organic with colors of green and brown … opening wider toward the end.” B6: “pattern in wall … honeycombed with thin lines of white and blue light.” (Supplementary Material C). The widening of A5 thus marks the transition to the panoramic B6 VF. The achromatic predominance of A-shapes likely reflects constricted VFs associated with loss of peripheral vision, as seen in hypotensive syncope (Lambert and Wood 1946), ocular pathologies (Edmeads 2003), and neurological disorders (Trauzettel-Klosinski 2010). Because peripheral color sensitivity is reduced, constricted VFs are perceived in grayscale (Hansen et al. 2009). This aligns with the retinotopic organization of the visual cortex, where the fovea is overrepresented relative to the periphery (Facco and Agrillo 2012a; Blackmore and Troscianko 1989). A conical VF emphasizes foveal brightness and peripheral darkness, consistent with NDE reports of dark tunnels with light at the end (Mobbs and Watt 2011; Parnia and Fenwick 2002). Reduced blood or oxygen supply further narrows the VF; under hypoxia, peripheral processing fails first, leading to tunnel vision (Leibowitz 1982). These findings suggest that A-type spaces may stem from retinal ischemia and cortical anoxia.

The achromatic dominance of A-type spaces aligns with the magnocellular visual pathway, which encodes motion and luminance but is color-insensitive (Livingstone and Hubel 1987). Under low illumination, vision shifts from cone-mediated photopic to rod-mediated scotopic mode. As rods remain active after cones cease, perception becomes essentially grayscale(Zele and Cao 2014). This rod predominance likely explains the characteristic achromaticity of A-type NDE spaces.

#### Categories of elliptic arcs VF boundaries (B1-12 and C1-4)

The VF boundaries in categories B1–12 and C1–4 are characterized by elliptical arcs subtending less than a full ellipse and showing strong chromatic bias. In our data, B shapes were predominantly chromatic, with 27 of 31 instances (87.1%) featuring color. Similarly, C1–4 shapes were mostly chromatic, with 13 of 20 cases (65%) versus 7 (35%) achromatic. C5 was treated separately due to its distinct VF boundaries, corresponding to a complete ellipse rather than elliptic arcs. In C5, chromatic bias was slightly lower than in C1–4, with 18 of 29 instances (62.1%) chromatic and 11 of 29 (37.9%) achromatic. The chromatic–achromatic distribution (Figure 3) shows clear contrasts among families: B-shapes display strong chromatic predominance (87.1% vs. 12.9%), C1–4 have higher achromatic proportions (35%), and C5 shows a similar ratio (37.9%). To illustrate these findings, we examine participants’ descriptions of B1–12 through both verbal reports and graphic representations. These reveal the chromatic qualities of the VF boundaries and show that, in participants’ graphic representations, B1–12 exhibit high curvature, while C1–4 display low curvature (almost flat).

Participants frequently described phenomena within B-type VF boundaries as vividly chromatic, with rich spectral variation. For example: “… a glass spectrum of all colors like a prism, like the Pink Floyd triangle with the rainbow spectrum” (Subject 1, Space 2, B2; Supplementary Material C; ESM, Graphic Representations, p. 5). “Many colors, especially greens and purple were vivid. Lavender, pink, magenta and deeper greens from bright to dark intensity! Very high clarity, more than on earth!” (Subject 17, Space 2, B10; Supplementary Material C; ESM, Graphic Representations, p. 84). “Life review … It appeared to be like microfilm moving through it very fast. It would pause on individual photos of myself and immediate family.” (Subject 10, Space 4, B7; Supplementary Material C; ESM, Graphic Representations, p. 46). “The room was made of pink and gold light and was porous…” (Subject 28, Space 3, B11; Supplementary Material C; ESM, Graphic Representations, p. 130). “It’s a vast open space… completely silent, and in my consciousness the color of that place where I sat is brown and orange like soil. I was looking at planet Earth.” (Subject 27, Space 1, B6; Supplementary Material C; ESM, Graphic Representations, p. 126).“… the ‘garden’ was huge with rolling hills, tall grass … vivid people walking and talking, children playing, animals roaming… There were the most wonderful flowers and trees in amazing colours.” (Subject 4, Space 2, B10; Supplementary Material C; ESM, Graphic Representations, p. 16).

These vivid descriptions reveal strong chromatic richness and a pronounced sense of spatial expansiveness consistently observed within B-type VF boundaries. Previously, we noted that rotations of S-L may influence the perceived orientation of the VFE boundary. In examining chromaticity, we further observed that the orientation of the VFE can also be influenced by the sudden appearance of salient elements, such as VMI (entities) arising within the VF boundaries. Such emergent imagery may bias visual attention and redirect the S-L orientation accordingly.

For instance, in Subject 21’s graphic representation of C5 (ESM, Graphic Representations, p. 101), the entity appears at approximately the 1 o’clock position within the encompassing ellipsoidal VF boundaries. In the subsequent representation of space B2 (ESM, Graphic Representations, p. 102), only the elliptic arc of the VF boundary onto which the entity is positioned is preserved. This selective attentional focus is further illustrated in space C3 (ESM, Graphic Representations, p. 103), where the S-L moves closer to the VF boundary at the location of the entity.

As a consequence of this S-M, the VF boundary expands into a broader elliptical arc VFE. A transformation is then seen in space C1 (ESM, Graphic Representations, p. 104), where both the entity and the VF boundary are shifted to the opposite side of the representation, suggesting that C1 is the symmetry of C3.

Finally, in the subsequent space, the C5 ellipsoidal VF complete enclosing boundary appears, but the entity is no longer depicted (ESM, Graphic Representations, p. 105).

These empirical observations support the hypothesis that attentional saliency and S-L orientation jointly modulate the perceived orientation and size of the elliptic VF arcs (B1–B12, C1–C4).

In contrast, A-type VF boundaries, defined by conical geometries, appear anchored in retinotopic coordinates (eye-centered, aligned with the body), whereas the elliptic VF arcs (B1–B12, C1–C4) and the C5 ellipsoidal VF boundary are represented in spatiotopic coordinates (world-centered, stable relative to external space). This distinction suggests that NDE VF boundaries may involve a shift from retinotopic (A1–12) to spatiotopic reference frames (B1–12, C1–4, C5), marking a transition from body-anchored to world-centered spatial organization.

Having detailed the chromatic richness of B-type VF spaces, we now turn to the C-type VF category. Compared to B shapes, overwhelmingly chromatic (87.1%) and rarely achromatic (12.9%) C shapes (C1–4, C5) showed reduced chromatic proportions (63.3%) and markedly higher achromatic reports (36.7%). The achromatic rate in C shapes was nearly three times higher than in B shapes, underscoring a clear perceptual contrast between VF categories. In participants’ space descriptions. Reports often refer to bright colorless light of extreme intensities across C1-4.“I was in an infinite space, transparent space, facing a brilliant source of light.” (Subject 36, Space 1, C2 Supplementary Material C and ESM, Graphic Representations: p. 157); “ … nobody, space was dark (back, left) and turning lighter (in front, right) turning lighter towards shape C5 (the light) light was a round circle, drifting towards the light drawn towards it like a magnet … when I got closer it was turning more a white light, the light was opaque, unified, circular” (Supplementary Material C and ESM, Graphic Representations: p. 139); “The light was it was a bright sunny day with love. I felt contained and embraced in the space … The light was pulsating and sparkling. It was retracting and expanding. It would pulsate randomly when talking. The tempo of the light was almost at the speed of a heartbeat.” (Subject 31, Space 1, C2 Supplementary Material C and ESM, Graphic Representations: p. 142). While C1–C4 were typically described as elliptic arcs often associated with bright but colorless or near-colorless light, C5 emerged as a distinct category characterized by a complete ellipsoidal VF boundary, marking a transition from partial arc segments to a full enclosing visuo-spatial configuration.

#### Categories of Ellipsoidals VF boundaries: C5

C5 is the only complete ellipsoidal VF boundary, unlike the partial arcs of B1–12 and C1–4. In our dataset, 18 of 29 cases (62.1%) were chromatic and 11 (37.9%) achromatic, showing lower color intensity than B-type spaces. Although its chromatic–achromatic proportions resemble C1–4 (Figure 6), C5 differs by its full ellipsoidal geometry.

Participants often described C5 as a 360*^◦^* enclosure of light that completely surrounded them, with greater luminosity than C1–4: “I was in the center of an immense bubble of light … it enveloped me completely, I can say that I was part of it … it was at the perfect temperature …” (Supplementary Material C). For Subject 40, C5 appeared as a state of ontological–affective fusion, echoing ego dissolution and phenomenological unity reported in altered states (Millière 2017; Fort et al. 2025): “One massive entity, brighter than anything I’ve ever experienced … I was connected … in total unity with it and everything else … It was like the Beatles line, ‘Limitless undying love, which shines around me like a million suns and draws me on and on across the Universe.’ It was achromatic and very physical.” (Supplementary Material C). Another described: “It was like pure white divine light behind the dark corridor, like behind a door; that room was like opening the sun.” (Supplementary Material C). C5 category may indicate a visual marker of ego dissolution where self–environment boundaries collapse (Stoliker et al. 2022).

Phenomenologically, the sense of immersion in C5 resembles the Ganzfeld phenomenon, in which the visual field becomes homogenously uniform. In C5, participants often described spatial extendedness as a space entirely filled with light. Consistent with Haun and Tononi (2019) account, conscious spatial experience can persist as an “extended canvas” even when perceptual content is empty or uniform.

## 5 Conclusion

This study presents a data-driven classification of NDE visual spaces into four geometric types. The identified categories: conical (A1–12), elliptic-arc (B1–12), low-curvature elliptic-arc (C1–4), and ellipsoidal (C5) each correspond to distinct VF boundaries and chromatic qualities. A-type conical spaces showing constricted, achromatic fields, whereas B- and C-types displayed expanded VFEs with chromatic dominance. B-types most strongly exhibited chromatic dominance. In contrast, C5 represented a volumetric, 360° ellipsoidal enclosure, forming a structurally distinct core NDE space.

Empirically, both elliptic-arc sets (B1–12, C1–4) may correspond to arcs of C5 ellipsoidal structure. Thus, B and C can be regarded as perceptual part of C5, while the A-type conical configuration appears at both onset and termination, directed toward or away from C5, suggesting that overall NDE structures reduce to two fundamental geometries: conical (A1–12) and ellipsoidal (C5).

A speculative interpretation proposes that B- and C-type arcs reflect patterned visual-field losses analogous to hemianopic defects. B-type configurations extend across more than one quadrant, whereas C-types typically present as a flattened hemifield boundary, most commonly in the superior visual field. Though not reducible to neuropathology, these recurrent quadrant-specific arcs imply structural parallels between subjective NDE reconstructions and known visual-field deficits.

## 6 Limitations and prospects

The main limitation concerns the use of freehand drawings to reconstruct NDE spatial structures. While providing essential first-person insight, their informal nature precludes standardized geometric analysis and quantitative measurement of shape, orientation, or scale. Future work will employ a 3D interface enabling participants to reconstruct experiences and standardized encoding of VF geometry, perceptual features and self spatial dynamics.

## Acknowledgments

I would like to express my sincere gratitude to Professor Roy Salomon for his invaluable assistance. I also thank Dr. Andrew Haun, Professor Ben Thompson, Professor Susana Martinez-Conde, and Professor Stephen Macknik for their insightful discussions. Special thanks go to all the NDErs, without whom this study would not have been possible.

## Supplementary Material

## A Plots And Tables

To keep the main body concise, this appendix presents several figures and tables from the Results section. Table 6 is the only table not discussed in the Results section; it provides the references corresponding to each OBE figure.

## B Classification of NDE Space Types (A, B, C)

For each NDE sequence, subjects selected the shape of the perceived space from a standardized catalog of pictograms (see Supplementary Material: *Catalog*). The catalog was divided into four families: A-shapes (A1–A12) representing conical spaces, B-shapes (B1–B12) representing elliptic arcs, C-shapes (C1–C4) representing flatter elliptic arcs, and C5, a complete ellipsoidal configuration. The C family thus included five total configurations, with C5 treated separately due to its complete 360*^◦^* enclosure. This pictogram catalog was created after Lerner et al. (2021), who demonstrated that NDErs naturally report these spatial configurations when describing their experiences. Using this catalog as reference, we defined four NDE space types (A1–A12; B1–B12; C1–C4; C5):

### Category A — Conical Spaces (A1–A12)

Conical VF boundaries defined by pairs of converging or parallel segments.

***A1*** Parallel vertical segments.

***A2*** Parallel horizontal segments.

***A3*** Parallel oblique segments (45*^◦^*), descending left to right.

***A4*** Parallel oblique segments (45*^◦^*), ascending left to right.

***A5*** Nearly vertical segments converging toward an apex above.

***A6*** Nearly horizontal segments converging toward an apex rightward.

***A7*** Oblique segments converging toward an apex above-left.

***A8*** Oblique segments converging toward an apex above-right.

***A9*** Nearly vertical segments converging toward an apex below.

***A10*** Nearly horizontal segments converging toward an apex leftward.

***A11*** Oblique segments converging toward an apex below-right.

***A12*** Oblique segments converging toward an apex below-left.

### Category B — Elliptic Arc Spaces (B1–B12)

Elliptic arcs (including circular arcs) defined by their angular extent (*≈* 90*^◦^*to *≈* 270*^◦^*) and orientation within the VF. The concavity and direction define the perceived enclosure of space.

***B1*** 90*^◦^* arc, opening lower-right, apex upper-left.

***B2*** 90*^◦^* arc, opening lower-left, apex upper-right.

***B3*** 90*^◦^* arc, opening upper-left, apex lower-right.

***B4*** 90*^◦^* arc, opening upper-right, apex lower-left.

***B5*** 180*^◦^* arc, concavity facing right; endpoints upper-left & lower-left.

***B6*** 180*^◦^* arc, concavity facing downward; endpoints upper-left & upper-right.

***B7*** 180*^◦^* arc, concavity facing left; endpoints upper-right & lower-right.

***B8*** 180*^◦^* arc, concavity facing upward; endpoints lower-left & lower-right.

***B9*** 270*^◦^* arc, opening lower-right; small gap at lower-right.

***B10*** 270*^◦^* arc, opening lower-left; small gap at lower-left.

***B11*** 270*^◦^* arc, opening upper-left; small gap at upper-left.

***B12*** 270*^◦^* arc, opening upper-right; small gap at upper-right.

### Category C — Flatter Elliptic Arc Spaces (C1–C4)

Arcs (angular extent *<* 30*^◦^*) with low-curvature, representing segments of large-radius ellipses or circles. Orientation defines the arc’s position within the VF.

### C5 — Complete Ellipsoidal Configuration

A full ellipsoidal (often described as 360*^◦^*) enclosing VF boundary, treated separately from C1–C4 due to complete enclosure.

## C Subject NDE sequence descriptions

### Subject 1 — Space 1 (B6)

“A backlight hill about 30 feet high seemed like dusk or evening eerie dark blue. There were stone stairs on the left in front of me and Jesus was toward the top wearing a white robe. I was a little bewildered slow emotion as he came toward me to meet me at the step and we had a discussion on the 3rd step we stopped It was a review or judgement of my life in this lifetime up to this point (I was about 6 years old at this time) So Jesus turned to me and said “You haven’t fully experienced life yet so we must stop here, however I want you to be aware that I will live and during life you may come into power and I needed to be aware that I will be judged when my time comes.”

### Subject 1 — Space 2 (B2)

“It was an orphanage It was a sunny day with some fluffy clouds like a normal scenery and then I saw a nun and went into a room with a bed the nun introduced me to 2 children who were relatives who had already passed, we were told to pick our musical instruments to play in Jesus’s garden. Vivid intense techno colors. Fluorescent, rainbow yellow, pink all the colors : a glass spectrum of all colors like a prism like the pink Floyd triangle with the rainbow spectrum. In space 2 motion of the first self-location is forward and left. “Vivid intense techno colors. Fluorescent, rainbow yellow, pink all the colors : a glass spectrum of all colors like a prism like the pink Floyd triangle with the rainbow spectrum”

### Subject 1 — Space 3 (C5)

“Light textures contrast dark like yin and yang black and white. It was like pure white divine light and that was behind the dark corridor like behind a door, so that room was like opening the sun.”

### Subject 1 — Space 4 (A5)

“The walls were perfectly parallel and perpendicular to watch others and it was a long corridor hallway. It was as if I was moving forward back … musical math very smooth with the exact same timing the hallway was going backward as I was moving forward at unison perfect timing.”

### Subject 3 — Space 1 (A1)

”I moved through a space similar to a hallway with no walls. As I looked forward, I was facing several beings (6-7). I don’t know if they were human, they appeared human in the form of the light they were emitting, they were glowing white. I sensed they were communicating with me, as I stared at them I began feeling their words. I could see my surroundings, but I couldn’t feel them. I didn’t feel the ground, but I didn’t feel like I was floating. I was feeling overwhelmed with all the new senses, that were not physical. The beings in front of me were remarkable, glowing very brightly, they seem happy and welcoming. No words were ever spoken, but they were not needed. I don’t know what messages I was sending them, but I feel them. Strangely, I was not scared, quite the opposite. I felt great! No body pains, no bodily feelings at all. I couldn’t feel my body, but there I was, I don’t remember ever seeing myself, i didn’t see my hands, my feet, nothing, but there I was. I was in wonder of my surroundings, what happened to my body? My thoughts were racing in all different directions. When I started picking up communication from the beings in front of me, it was a little overwhelming. I was trying to process their feelings at the same time I was asking myself “why can I feel them?” I was trying to understand logically what was happening, telling myself ‘this is impossible’ Once I let my self connect with them, all their feelings quickly turned to comfort and love. That word does not describe it well enough. Unconditional love, love for a child or favorite pet, it’s the kind of love without boundaries. They were telling me I was safe. It was going to be fine. They were there for me. They will answer all my questions. They are there to help me understand. They were welcoming me. They said I was home. I felt myself move closer, as if I was meeting my family at the airport after you’ve been away from them for months. The love between the beings and me was growing as we moved closer, like we were connecting more. The air around us was glowing. I realized they were all communicating with me, it wasn’t coming from just the person in front. They felt like family, even if they weren’t my family, it felt like we were one. Like we were all ONE. Then, suddenly, I heard Cindy yell my name. She sounded scared. She yelled my name again, louder. And again. She seemed very upset. I paused and knew I had to go back. I couldn’t leave her alone. As soon as I had that thought, I felt myself move further away from my new friends. I felt like I was moving to my right, as if I were slowly being pulled by a rope. The tunnel vision returned and it went black. I started to wake up. It was like waking up from surgery, very groggy, I had no idea where I was or what happened. I was back in my body. Laying on the ground with my wife and paramedics looking at me in suprise.”

### Subject 3 — Space 2 (B11)

“As I looked forward, I was facing several beings (4-7). I don’t know if they were human, they appeared human in the form of the light they were emitting, they were glowing white. I sensed they were communicating with me, as I starred at them I began feeling their words. They were communicating with me through thoughts and feelings. Once I felt them, it all came very fast. It felt like a long minute, collecting myself in my new surroundings before I noticed them and then noticed them communicating. I could see my surroundings, but I couldn’t feel them. I didn’t feel the ground, but I didn’t feel like I was floating. I was feeling overwhelmed with all the new senses, that were not physical. The beings in front of me were remarkable, glowing very brightly, they seem happy and welcoming. No words were ever spoken, but they were not needed. I don’t know what messages I was sending them. Strangely, I was not scared, quite the opposite. I felt great! No body pains, no bodily feelings at all. I couldn’t feel my body, but there I was, I don’t remember ever seeing myself, seeing my hands, my feet, nothing, but there I was. I was in wonder of my surroundings, what happened to my body? Why can’t I feel my body? I was curious, excited, optimistic, I felt great. No aches, no pains. My thoughts were racing in all different directions.”

### Subject 4 — Space 1 (A1)

“dark to start, as I started to travel along I saw images on the walls of my life (life review) in vivid colour with sound, like a movie, good resolution. I was able to slow to watch. images were good and bad. became lighter as I traveled on to the end.”

### Subject 4 — Space 2 (B10)

“The being at the end was pure light and only saw a shoulder and down. the ‘garden’ was huge with rolling hills tall grass. there were very vivid people walking and talking, children playing, animals roaming. I could hear the most wonderful music. There were the most wonderful flowers and trees in amazing colours. everything was vibrant and felt warm and welcoming. I felt that I wanted to enter but the being told me that it wasn’t my time and that I had to go back.”

### Subject 5 — Space 1 (A1)

“I entered this corridor, with cabins on my right. On my left, there was nothing. From inside the cabins came a chant, I heard men studying (the Torah) and it was melodious. A voice in my ear whispered to me: Avraham Avinu, Rabbi Akiva. I was extremely happy to be there, in my place, I moved forward with confidence, I saw the cabin doors closed, I just heard the melody. All the colors around me were rather dark, gray, black of different shades, but not in a negative way, more like something hidden, not visible. I moved forward feeling completely serene and confident, curious too.”

### Subject 5 — Space 2 (C5)

“I was in the center of an immense bubble of light, I saw no edge of this bubble, it enveloped me completely, I can say that I was part of it, this light was the most beautiful ever seen, it was at the perfect temperature, I was wrapped like in silk, like in a down but the lightest in the world, like in cotton wool, I was in my place, where I always dreamed of being without the namely, I felt the greatest joy, serenity, deep calm and the greatest connivance with myself and with G. Completely in my place. enveloped and part of this light, I was not outside it. I didn’t want to move anymore.”

### Subject 6 — Space 1 (A1)

“I encountered my Spirit Guide. She was already there waiting for me. She was wearing a dress, with dark hair and we were having an ongoing conversation telepathically. The space was silent and black with shimmering velvety coppers, silvers and gold.”

### Subject 6 — Space 2 (C5)

“The space was cottony white all around and appeared to have no defined walls or ceiling or floor. There were many family and friends there greeting me. I was able to touch them and we shared memories when we touched. I could remember my memories as if I was reliving them and I could remember from their point of view as well. I couldn’t hear anything other than voices. I couldn’t smell anything. A round portal opened in the white when I thought of my family on Earth and I could see them. I felt a growing sense of love and peace. When I remembered my past lives or had a question I felt an overwhelming sense of standing in an electric stream of knowledge that flowed through me and I instantly knew the answer to my question or went to a space where I learned the answer. The flow of knowledge was beautiful and rainbow colored in vibrant colors. It felt like Home to me, that it is who we really are and we are it expressing itself. The knowledge was omniscient.”

### Subject 6 — Space 3 (B12)

“I entered a space that looked like outer space and felt infinitely big. I felt a sense of movement forward and backwards and at single u-turn. I was moving at incredible speed so that the stars flew by like in the Star Wars movies when they are at light speed. I came to a planet and zoomed in on one being that I instantly recognized as myself. I then moved backward off the planet and zoomed forward again. I passed through a solid black space very quickly, instantaneously, and then very quickly arrived at a second planet where I saw another being that I recognized as myself. I then moved backward off the planet and then made a u-turn and returned to the white room. I didn’t hear anything in this space. I spoke telepathically with the two beings I met. I didn’t feel any type of temperature.”

### Subject 6 — Space 4 (C1)

“I left the white room and entered a space which was like outer space where stayed in one space and was able to see all of the past/present/futures from my current lifetime on Earth. It was like watching a movie but I could feel the emotions of everyone in an overarching sense-so if it was the future of Earth then I felt the emotions of the groups of people on Earth being affected in that scenario.”

### Subject 6 — Space 5 (C2)

“I encountered humans as groups of people as I saw the past, present, and future of Earth. It was very clear and colorful. It was more real than seeing with my human eyes. I could also feel the emotions of the humans that I saw. I could use all of my human senses, but with more depth and clarity. It was the best of the best as far as feeling, and seeing, and hearing, and all of the senses. The whole Earth was communicating with me as One. Plants and animals were vividly alive and I could see that they are One with humans, and each is just as important as anything else on Earth. Being a vegan, for example, does not rectify the slaughter of an animal because plants are just as full of the Life of The Other Side as every other being on Earth.”

### Subject 6 — Space 6 (C5)

“I encountered my family again and my life guide. It was all very, very, vivid and colorful. I had a life review wherein I saw highlights from life of good and not so good things I had done. I learned that no one judges me, but me. I was very disappointed with myself even though the things I thought I had done that were negative were not very bad by Earthly standards. However, I was viewing my human choices from my angelic self’s viewpoint. As I was feeling disappointed I felt all the other beings soothing me and reminding me of how hard it is to live on Earth. They were filling me with their love and their lack of judgement was overwhelmingly lifting me out of my disappointment. We interacted with each other via telepathy.”

### Subject 7 — Space 1 (A1)

“I was contained in a tubelike space and it was invisible. I could see normally around myself.”

### Subject 7 — Space 2 (C2)

“At first the space felt tubular. It expanded and became wider as it progressed and became C2. I could see infinity. It appeared in the distance like space. Black with white flecks like starts and a grey haziness around. It was infinity opening in all directions at once.”

### Subject 8 — Space 1 (A2)

“There were many tiny specs of light, that I related to being other humans/souls traveling upwards, just as me. They were located all around me, at many distance, but all were quite far away from me. Also I encountered (in my thoughts) a god-entity that I had a negotiation in my thoughts. He shared with me understanding of the inter-connectedness of everything and how “all is one”. The space that this happened in, was vast, dark, open around me, with a ceiling with a hole (like a man-hole) in it. I was not able to move, I still felt constricted in fetal position, but could look 360 degrees in all directions. I could still see my physical body in our bed with my wife and our children.”

### Subject 9 — Space 1 (A5)

no description

### Subject 9 — Space 2 (A5)

no description

### Subject 10 — Space 1 (A5)

“moved very fast. walls seemed to be organic and had colors of green and browns but also had tones of what appeared to be veins. it appeared to open wider as I moved to the end of the tunnel”

### Subject 10 — Space 2 (B6)

“noticed pattern in wall. Honey combed with thin lines of white and blue light that would fade in and out. it followed the pattern of the honey comb. To the left there are three galaxies that are 2 on top and one on the bottom i tried to bounce the lower galaxy in my left hand it would move up and down hovering above my hand.”

### Subject 10 — Space 3 (C5)

“Still felt contained, but much larger area. God force entered from in front of me toward the right. I had the feeling of wanting to lift my head up but could not or felt that I shouldn’t. Was given huge download of information ( a knowing). Felt incredible amount of love like I have never felt before complete forgiveness and acceptance. Messages are telepathic but so strong as if its a voice. was very specific on some topics general on others. Full knowledge of everything. feeling of complete exposure of secrets forgiven. My location was dead in the center of this space when this was all taking place. there was a back drop noise of many people talking as if in a large venue. Could pick up bits and pieces of what they were saying. But the god’s voice was very direct and clear.”

### Subject 10 — Space 4 (B7)

“Life review. Looked to be consistent of caramel in a thick liquid form. It was in a ribbon formation that seemed to float in the space to my right but floating and moving. As the ribbon moved closer to my vision I was able to detect movement in it. It appeared to be like microfilm moving through it very fast. It would pause on individual photos of myself and immediate family. These are very happy memories of times with my family. Didn’t seem to be of any one occasion that I can remember. My overall feeling was one of being content. I had no fear of punishment was completely accepting.”

### Subject 10 — Space 5 (A9)

“Started my return to my body. Started out as a feeling of descending and then fast change to an ascending motion again this tunnel was different there was a light at the end of this one it was dim not a brilliant light. I then emerged in the ICU just for a couple of seconds and then felt as though I was thrown back into my body feet first.”

### Subject 11 — Space 1 (A5)

Translation in English:“A tunnel with a light at the end; one moves very quickly, with a sensation of speed (touch).”

### Subject 11 — Space 2 (C2)

Translation in English:”Like a movie screen, one reviews their life along with the emotions it evokes.”

### Subject 11 — Space 3 (C5)

Translation in English:”One is free, fully oneself, without age or gender, one has finally come home, and time is not experienced. One sees everywhere at once.”

### Subject 11 — Space 4 (B6)

Translation in English:””One has just instantly traveled an astronomical distance, and has arrived; one appreciates the place like a child — it is colorful, magnificent, very vast, each place is unique, the color of fire: red, orange, yellow, quite vivid, like clouds. It is hard to describe the texture; it is very beautiful and peaceful.”

### Subject 11 — Space 5 (C5)

Translation in English:”One has just instantly traveled an astronomical distance, and has arrived; one appreciates the place like a child — it is colorful, magnificent, very vast, each place is unique, the color of fire: red, orange, yellow, quite vivid, like clouds. It is hard to describe the texture; it is very beautiful and peaceful.”

### Subject 11 — Space 6 (B6)

Translation in English:“During this journey, it is as if one zooms toward a destination and suddenly finds oneself there almost instantly. It is a succession of C5 and B6; when one arrives, one slows down. During all three displacements the sensation is the same: one chooses the destination, zooms in on it, arrives while slowing down, and appreciates the place. On one of the journeys, one brushed past a place where one should not go — it is sad, dark, and frightening; if one goes there, one is destroyed. It was very, very vast. One must not zoom in on that. It is very dangerous. Each time one jumps from one place to another, the horizon line is narrowed like B6, and at each stop one returns to C5. The field of vision changes when one moves, or when one is at rest.”

### Subject 11 — Space 7 (C5)

Translation in English:”Here, we are motionless, spinning on ourselves, and the Earth comes into our field of vision.”

### Subject 11 — Space 8 (B8)

Translation in English:”Here, it is the dizzying fall toward the Earth and the body; the visual field is directed downward and very narrow, and one does not want to return to Earth.”

### Subject 12 — Space 1 (A5)

“Dark at the beginning. At the end of the corridor the light was coming through.”

### Subject 12 — Space 2 (B9)

“In the big space opening into the corridor was where the light came through. The opening corresponded to the light coming through. The pure light was shinning and giving love and bliss to me and to my infant son. I could feel the love emanating from the light. My infant son was beaming with love and bliss. He was shining in the light as he floated off into it. The last image I remember was that he had a blissful smile on his face as he floated away from me towards the light.”

### Subject 12 — Space 3 (C2)

“The light was lessening. As the light was diminishing, my feelings turned negative; I realized I didn’t want to surrender my son to the light. I had the thought that I wanted to get him back. No, I did not have the sensation of moving in this space. No motion.”

### Subject 13 — Space 1 (A5)

“Felt I was going though star fields; black. Saw stars that were bright but they were stationery; I was moving. I did not felt cold.”

### Subject 13 — Space 2 (B6)

“Bright light yellow walls, grandfather was well and looked happy and delighted to see me. glowing with light, tan pathway into courtyard, beautiful, vision of paradise, colorful colors, rainbow colors, music, beautiful music melodic, blue a gentle sound, pink fragrant flowers, color matching with sounds. The courtyard was beautiful architecture, pillars, sand in color, with beautiful flowers and beautiful fountains making lovely sounds. The most beautiful flowers I have ever seen.”

### Subject 14 — Space 1 (A5)

no description

### Subject 14 — Space 2 (B5)

“Within the space, it was filled with bright light, and I felt like the light was the being. I felt like I should turn to face it, and the light was brighter and more intense, and loving.”

### Subject 15 — Space 1 (A5)

“it’s look like night time (little dark),space is like I’m walking in water”

### Subject 15 — Space 2 (B6)

“The space is like semi sphere from my view point. it has some colors and light but there is no sun. I can hear air sound.”

### Subject 16 — Space 1 (A)

”Field of vision was infinite, a tranquil, dark energy line. Outside it were many frantic, assorted rainbow-color electric impulses coming toward the big, dark line but not effecting the big line. Wherever I move, the big line followed my perspective (up down left right…). Turned around for A9, A10, A 11, A12. Intense, pleasant aroma (a cross between amnionic fluid, brewer’s yeast, sweet electric burning, Fenugreek, & St. John’s wort.). Auditory: I heard in harmony. The sensation was as if I were part of the dark matter and it was fluid and calm. A vivid rainbow-colored ““angel ray”” like a thin, single, arched sunbeam extending upward.”

### Subject 17 — Space 1 (A7)

“I saw at end of tunnel a bright white light!”

### Subject 17 — Space 2 (B10)

”colors were external but within like a backlight. Many colors, especially greens and purple were vivid. Lavender, pink, magenta and deeper greens from bright to dark intensity! Very high clarity, more than on earth! texture of the grass was like velvet, the petals of flowers were like soft, transparent, satin texture! I saw my aunts Elizabeth and Linnie as they were when they were younger women-thirties maybe. i only knew them when they were sixty-seventy years old! I am seeing my maternal grandmother, Ida Mae, as a young woman! I never knew her alive as she died before I was born by many years! I saw a man to my left. No name was known, but I seemed to know him as Peter, he was gardening the gate leading into the green pastures and flower gardens. He asked me a question: Vicky what have you done with your life my heavenly Father gave you? Immediately began to cry.”

### Subject 17 — Space 3 (C5)

“No words exist to describe this existence! All knowing, separate from the light! I wanted to die, but I was begging the light to release me to my body so I could take care of my two little boys, I wanted to stay in the wonderful light, but I knew I had to make changes in my life! I had to go back to earth!”

### Subject 18 — Space 3 (A8)

“I first saw all of the broken pieces of wood and cement and I remember seeing a mattress - it was a mess, everything was everywhere. But as I kept moving through the space it turned into more of a tunnel with just the bright yellow and white light. An angel came and she took me through the tunnel. But her form was not very defined. But I knew it was her because I had called on her for help. A man had told me that she was my guardian angel a couple weeks prior, and to call on her when I needed her. So when I woke up buried in the landslide, I called on her. And then she came and took me through the tunnel. But when I came out of the hole at the end of the tunnel she was not there with me. But it was like an actual rebirth, as I came out of the hole in the ground naked (because I had been sleeping naked). And I came back into my body and awareness at that moment.”

### Subject 18 — Space 3 (B6)

“It was if I was watching a screen, a movie of all the moments of my life flashing by one by one. I saw them all individually, but also kind of in a blur. And then I saw all of the people I’ve ever met in my life and had some kind of interaction/connection with, even if it was just for 1 minute. From my family and friends to even people I’ve met on the street or someone who checked me out at the grocery store. And I saw those people in the moment that I met them and I remember the moment of connection - specifically their eyes looking at me.”

### Subject 18 — Space 3 (C5)

“At this point, it was as if I was shown the secret to life. And I remember being in a completely open space, perhaps even floating in some sense with a bright light all around. Maybe even some kind of pastel rainbow colors - nothing too bright. All the lights and textures were moving around a lot. And it was if they pulled everything away to reveal the simplicity of life at its core. And I had this profound understanding that all of this “stuff” in our lives is actually an illusion and it doesn’t matter. That it is complicating everything. That the secret to life is actually very simple. I was in complete awe and kept whispering, “don’t ever forget this”.

### Subject 19 — Space 1 (A8)

“in A8, I was aware of others moving outside to the left like particles of light B4, pastel, gold, glimmer, soft, cloudy rainbow.”

### Subject 20 — Space 1 (A9)

no description

### Subject 20 — Space 2 (B6)

no description

### Subject 20 — Space 3 (C2)

no description

### Subject 20 — Space 4 (C5)

no description

### Subject 21 — Space 1 (A9)

“Escalator in a pure black space. Floating upwards through pitch blackness. 45 degrees, constant motion.”

### Subject 21 — Space 2 (C5)

“The landscape was of nature, rolling hills, and wild grass. Everything felt alive. The grass was glowing a white golden light. If you zoomed your focus on the grass, you could see into the light within it. If you did not zoom into the grass, you would just see glowing grass. There was a full spectrum of color. I was standing in the middle of a 360 degree sphere with a white, milky border. I encountered a being which was right on the edge of the border within my attention. He looked to be a man in his early 30’s, dressed in normal clothing. As we made contact I could sense that he was not human. The entity first appear as the perfect human as it came closer it kind of morphed a little bit more a light fluid being entity … telepathic communication with the being”

### Subject 21 — Space 3 (B2)

“The being from a distance appeared to be human, male, 33 years old, 6 feet tall, broad shoulders, fit/athletic build, short brown hair, 175lbs. Looked like a male model aesthetically. The border of the space was organic. As if it was a white moving organic substance between two panes of glass. The surroundings were still of nature, rolling hills, and the being was sitting on a large rock.”

### Subject 21 — Space 4 (C3)

“I encountered the being who was now very close to me. He was very clear and human-like at a distance. Now he appeared somewhat distorted. His shape became more translucent, and fluid.”

### Subject 21 — Space 5 (C1)

“The being is now on my left, at exactly a mirrored image to the prior position. In the prior position he was at 1 o’clock, now he was at 11 o’clock. My life review happened from this position. It felt like a filtered hologram that became very vivid if I put my attention into it. The feelings were of my experiences and the experiences of others. It also entailed decision making in my life as well as opportunities to change.”

### Subject 21 — Space 6 (C5)

“There was a growing light that grew to the point that I was encompassed in it. I felt fused with it, as if I was a part of it. It was a glowing white, silver, and gold. It emitted a feeling of love, that I was love, and a part of it. A feeling of being touched by grace.”

### Subject 22 — Space 1 (A9)

“Moved trough tunnel, started to lighten as I progressed, started black as night finished as gray early dawn.”

### Subject 22 — Space 2 (C5)

“Like a babe in its mother’s arms…became light, Jesus at my right, bearded robed, there to show me the way out. At that time heard/felt motion/noise behind me, asked what it was, told that I was still loved and could return or merse on …saw the sounds and colors a guide appeared at my right shoulder, stated he was there to lead me to next experience (guide was Jesus/love, communicated without words, after seeing lights and sounds associated with my father I asked what they signified, informed that I was still loved and will return if I desired, next experience was awaking from coma.”

### Subject 23 — Space 1 (A9)

“Loud buzzing and vibrating sensation when I my vision started to tunnel. Enveloped with an organic tunnel that was completely black, but had iridescent qualities.”

### Subject 23 — Space 2 (C5)

“Space felt tubal with no sense of an ending, but not claustrophobic.”

### Subject 23 — Space 3 (A8)

“The space felt like it was getting smaller and darker as I was being pulled backwards into it.”

### Subject 24 — Space 1 (B1)

“I was a gorgeous aura swirling bright pink going anti clock wise I was almost vibrating i felt guided i knew my deceased grandfather was guiding me without words “its not your time” but he never had to say it… i just knew it. back to earth i went but before i went back to body i heard my boyfriends thoughts saying “what am I going to tell her family” over and over again. and when i asked him about it days later he said yes he was thinking that. I was telepathic. It was the most beautiful experience of my life.”

### Subject 25 — Space 1 (B12)

”A big voice, he was masculine and commanding; a sovereignty; no discussion but unconditionally loving. Oily environment seaweed like dense. “closing on you” the sensation of touch is not on the physical body but on the self-location.”

### Subject 25 — Space 2 (C5)

“I felt the big void, oily and thick blackness. The emotion was so intense, the Presence that entered gave the void a limit or structure. The Presence also pushed back the feeling of void and extreme loneliness and fear. As if holding back the oily disgusting negative torment.”

### Subject 25 — Space 3 (B10)

“I see the negativity swirling around me and the Voice is to my right and feels enormously big, He said, “If you do not forgive this is where you will stay.” I argued with Him saying, “I’m a Holy Spirit Christian what am I doing here?!” The Voice said again, more authoritatively, “If you do not forgive this is where you will stay.” I remember vividly making a choice in my heart of hearts, the very center in my being that I would commit to forgive. Even though I had no idea who I was forgiving, yet I knew it was for the future of my life on earth in my body. Then immediately upon deciding, choosing to make a commitment to forgive that the bottom of the sphere opened and I looked down. I didn’t know to look down but it was as if the Presence, He told me to look down. I saw and felt all of the negativity swirling down toward the bottom of where my feet should be. I saw the black thick, oily substance going down a funnel and then I felt like i was being vacuumed, or sucked down the tunnel. I remember feeling frightened but the Presence assured my that I was safe and then I felt more of the vacuum, very fast and so, so fast into the tunnel. Then I remember crashing into my body and then my eyes opened.”

### Subject 25 — Space 4 (A9)

“The size of the sphere was big, then the bottom opened and the swirling started to funnel down. The opening got smaller and smaller as I was vacuumed down into the funnel. It felt almost too small but I went through it and crashed into my body.”

### Subject 26 — Space 1 (B4)

“Prior to the syncope/NDE, I was leaving the Alvord Desert and had been traveling on the road at this point for roughly two weeks, with the weekend break in the alvord desert to film for discovery channel. Upon leaving the alvord desert, I was filled with excitement for a short time and as the drive continued, along this two way road with the desert surrounding me, due to the drive creating long scape of seeing the same thing and a mirage appearing in the distance in front of me, i suddenly became sleepy. I remember slapping my right cheek a few times trying to wake me up and that’s the last thing i remember as the syncope occurred shortly there after. The space I entered was shaped as one could imagine what an infinite space would be. There were no confining walls that were defined but this airy light pink clouds that surrounded me almost as if it were a dome. I could see the texture of these ‘clouds,’ and resemble what you would see in the day sky clouds, with semi transparency, that blended together like an oil painting. I was walking in a straight line, pace was steady and slow, looking to my right and left taking all of it in, then suddenly something grabbed my shirt, bunched in their hand in the place where your heart is located (middle of my shirt) and i was forcibly tossed from this infinite realm and back into my present body in my car. In this time my car had gone off road and i awoken to just a mere inches in front of a speed limit sign and a sudden jerk of the wheel brought me back to the road. This was where i could process what i had seen in the moments of being tossed. I was able to grasp a glimpse of what appeared as an angel, these exquisite white wings, the feathers incredibly detailed and layered onto one another and his face was that of a Greek god, very symmetrical and with polished hair like you see in Greek statues. His hair was blonde and his skin was white. His features would fit the ideal beauty we would find in someone, the light was coming from within his body, and his wings were pearly white, hair an ashy blonde, and skin glowing white but not ghostly, just smooth and angelic. His figure was holographic, 3D. There was a telepathic communication with my angel that I was in a place of what would be where my spirit will first go after i leave my earthly body, it was telepathically said “it wasn’t my time.”

### Subject 27 — Space 1 (B6)

“Its a vast open space, I was sitting in a place, there are no other beings, Its completely silent and in my consciousness the color of that place where I sat is in brown and orange like soil. I was looking at planet earth. The Planet Earth was in beautiful Blue and Green and I was thinking about people sitting there. Like away from whats happening and seeing it as an observer. There was very dim light on the planet where I sat, but there was good and bright light on earth. I was in a calm silent state but felt like watching people moving doing there normal daily activities. Its was like watching Google Earth from a far off planet. I was in a sitting position, sitting directly on some rocky formation of 3-4 feet.”

### Subject 28 — Space 1 (B6)

“started human size and became infinitely big, it expanded; I went down toward the underwater pool light, but it was not an ordinary light, it poured out with light toward me and expanding inward to infinite depth. The light was as if breaking apart into tiny particles, no edges, like dust motes in sunlight all around the edges. At first I was separated and looking at the light, moving toward it, then I went into it.”

### Subject 28 — Space 2 (C2)

“I was in an open space, like a vast landscape, very beautiful, everything beautiful and luminous: most vividly, I felt like I was home–in a home that I had forgotten I had. I felt I belonged completely, many beings (like people) were generally around and they all knew me and I felt completely and totally loved and recognized. The beings did not have definite edges, they were more blurry. I was in motion the entire time, as if running or moving quickly and playfully across the landscape. Ahead of me, the landscape changed and there were perhaps buildings or something like buildings.”

### Subject 28 — Space 3 (B11)

“I was in a room made of light with beings that were also made of light. We communicated telepathically, without words. The room was made of pink and gold light and was porous, like light pouring off and through everything. I was with four or five beings, I have the sense that one came through the back area of the room to join us and stand in the back. I have the sense that the beings could take form like humans if I needed them to, but their essential selves did not have a specific physical/visual form in the way we usually perceive people. They were very kind and gentle and lovely toward me and encouraged me gently to return to earth, saying first, “do you want to go back” and then “won’t you miss your mommy?” and then “what about your dad?” But I wanted to stay there and did not want to go back to my earthly life. They told me my father (who was responsible for my drowning) would feel pain his whole life if I didn’t return. They let me feel what his pain would feel like. It was like having a sharp, broken piece of glass stuck in my heart. I felt the pain for myself. Because I still didn’t want to go back, they assured me he would feel that pain his entire life, even when he was an old man, if I didn’t return to my earthly life. I felt that was wrong, that no one should have to feel that much pain, even my dad. So I agreed to return.”

### Subject 29 — Space 1 (B9)

“No other beings”

### Subject 29 — Space 2 (B11)

“No being within that space, just observing the scene within that space that had the real physical people of 2 friends and my girlfriend”

### Subject 29 — Space 3 (B12)

“Only observing the characters in the scene unfolding below.”

### Subject 30 — Space 1 (C2)

“Nobody, space was dark (back, left) and turning lighter (in front, right) turning lighter towards shape C5 (the light) light was a round circle, drifting towards the light drawn towards it like a magnet. colors left dark blue - toward right - turning lighter blue eventually yellow/white (light) like color between the sun and a full moon - when I went towards it it was a powerful golden light, when I got closer it was turning more a white light, the light was opaque, unified, circular”

### Subject 31 — Space 1 (C2)

“Membrane that you could see through it but I couldn’t cross it. I was part of the space. The light was a bright sunny day with love. I felt contained and embraced in the space. The light was on the left in place of the face. It was sparkling and radiating love and calm. The light was the body and the head except for the hand. The light was pulsating and sparkling. It was retracting and expanding. It would pulsate randomly when talking. The tempo of the light was almost at the speed of a heartbeat. The voice was in my head by the light and the person on the other side of the veil/membrane. It was an older lady who knew me but I did not know her.”

### Subject 32 — Space 1 (A9)

“The space was dark, I was not aware where I was until everything came to me in an instant shot, It was like a vortex in out in-twinned like I was all parts of existence in the single moment, There was awareness from all the visuals of the universe existence and creation of how everything is in his world and beyond what this world is, then into the shot of my life review”

### Subject 32 — Space 2 (C2)

“started at 3 clock wise, was coming the tunnel of dark and as I was at the top of the light or in the pace of light I had seen starting from 6 o’clock My present life overview of my children their life, from 6 o clock I saw every path of my journey, even other parts I have not lived or choices i could make, 9 o’clock I saw people who I will meet, Faces of specific people who I will help and what choices I should take all was shown to me what I am still needing to do”

### Subject 33 — Space 1 (C2)

“Mountainous, high up, going downhill. Muted colours yellows, oranges through trees, in remote bushland. Like a winter’s day. I could feel the leaves on my hands and knees as I went down, there was a river at the bottom. I heard voices but it didn’t feel like meeting a ‘being’ more like memories or a static-y radio signal.”

### Subject 34 — Space 1 (C2)

”It was like a matrix, with many, many grid points, all connecting to each other in multiple dimensions. I felt that if I entered the matrix, I would be able to travel anywhere in the entire universe, simply by thinking about it. This was because everything was connected. Colors: black, but not dark, if that makes sense? I don’t recall feeling any textures. It was completely peaceful. I knew that everything had a purpose, nothing was at conflict with anything else. I don’t believe this in real life (that everything has a purpose), but I believed it in this space! It was so, so, happy and calm. I had the sense that my son’s spirit was close. Then I realized that this was the passage between birth and death, and these two things are simply the same process, but in reverse. One is spirits coming into the world, the other is spirits leaving the world. I knew death was nothing to fear. And I knew my son was entering the world through this doorway.”

### Subject 35 — Space 1 (C2)

“First ordinary people that I didn’t know. Then I saw Super natural, powerful, many more colors than we have, penetrating the and exposing them like pure crystal, completely transparent, so I could read their mind and see their whole life story. They all fell forward and together said the same prayer. Terrifyingly awesome, Holy, Holy, Holy, Lord of Lords and King of Kings, Majestic God of all creation. I am a vile reprehensible sinner, unworthy of your presence, worthy only of death and hell. Then they all got up, turned left and ran away from the light and to my left sa fast as they could. As turned left to see where they were going I saw the gates of hell open and the running in as fast as possible to get away from the light. I saw inside was dark, orange and very hot, like molten lava, but mostly sensed hate and shame.”

### Subject 36 — Space 1 (C2)

“I was in an infinite space, transparent space, facing a brilliant source of light. The light was incandescent, the brightest light I could imagine, and I was wondering how something was so bright, yet I could still look directly at it. I was floating, very slowly toward the light. It was silent and felt very peaceful. I had no worries or concerns about whether I was going to live or doe, I just wanted to move toward the light. In front (and slightly below, to the right) was a sort of black solid shape, shaped exactly like a backwards #7, and inside that shape was complete blackness. It was like a counter or solid shape that I had to get around in order to get closer, to continue moving. I felt this was an impediment to me getting toward the light, and I had to move around it. I looked up, and to my right, and saw my parents and my beloved aunt, they were leaning on their elbows, looking over a shelf, down toward where I knew my physical body was in the emergency room, far below us. I was very curious about getting closer to the light, and was floating toward it, although I had no control over my movement, I was drawn as if by a magnet. I was comforted by the presence of my family members, and was glad that they were watching what was happening to my physical body. As I got closer to the lighted area, I felt a very authoritative voice say to me. I did not get past the black space. “This is not your time, you are not going to die today.” And then in an instant, I was back in the mayhem that was happening in the emergency room.”

### Subject 37 — Space 1 (C3)

“All around me (extracorporeal self) was hazy-milky. It was some 70 times size of a human. I could see the body lying flat covered with two blankets and surrounded by three heaters and doctors shouting ‘body ko heat do, body ko heat do’. There were six doctors. Two on either side of my body holding surgical equipments (a female doctor on my right), one on my feet side looking at the machines, one near my neck (right side) checking the blood infusion. On the floor (right side of my body)I could see a can full of blood and a metal tray full of blood clots. A nurse inside the operation theater behind the doctors at my right hand side and one nurse breaking the news of my death to my family outside the theater. There is a small gallery outside the theater where my family is standing (my husband, mother, mother in law, husband’s friend and two unknown people. Everyone crying except my mother who was talking to the nurse. On right side of my body was blood being infused in the body and on left side near to wall was saline administered on my left hand. A table was lying some four feet away my head. There was a door to some room or store on the right hand side. There was blood soaked gauge kept in one metal tray on the small table on my right side.”

### Subject 38 — Space 1 (C4)

“I encountered floating spheres glowing blue and white. They were gently floating around me, fading in and out of view. They did not seem to be moving.”

### Subject 39 — Space 1 (C5)

”I had a feeling of being my own energy field and being part of the whole space,. the border shape felt like electrical waves. I could see the blue sky around me like high resolution, there were a few cumulous puffy white clouds in the distance. I think the clouds were moving very slowly and gracefully going past me at a linear but clockwise direction. I was alone however I was aware that if I want support there were beings out of my view but waiting to see if I would call on them, I also felt/sensed that i was in a telepathic communication like a negotiation of deciding to return. to this day I have a sense that while I made the decision to return if I had not, I may have experienced more negotiation communication before leaving to go ‘home’. I did have a strong sense of home not being earth. the space I was in was a holding space to decide which direction to go, and I realized that to go’ home’ was just a thought with an intention away. but home was somewhere deeper into space. I was aware of the energetic self (C5) that it was like translucent also transparent as i could look through it, just as if I was part of the atmosphere. I was able to look down at the red roof of the backyard garage and the concrete driveway where my body lay and the blackboard chalk board where i had done a drawing that was attached to the fence boundary of where I lived as I considered my option of ‘home’ in space or earth, i recall looking towards earth and with the thought to return to earth I became aware of an umbilical chord type of connection shining like fine silver I just had a knowing that it was connecting my spirit self to my physical self and I knew that I was still alive and breathing as long as that cord remained attached. the decision to go ‘home’ would have torn that connection cord, I have a very clear memory of being annoyed at my mother and I did not want to depart this earth with this emotional connection - unresolved. even though I was only 7 I had a very clear awareness that I did not want to leave this life with attachments of karma (but i did not know that word at that time) I just knew that when I leave and go home, i will not have any links to other people I will be free to be, individual me. therefore I decided to return and in that decision the next vision was my feet going first like I was skidding back into life(resistant, however I knew I had made this decision myself, even if I didn’t return with enthusiasm. next I was hovering at ceiling height in the room where my body was, just before returning to my body.”

### Subject 40 — Space 1 (C5)

“1 massive entity, brighter than anything I’ve ever experienced. It sparkled and pulsed in a living/organic way. the most intense light I’ve ever seen. I was connected… in total unity with it and everything else. I had no need to speak because I could/did know anything in a penetrating way. Penetrating in a 100

### Subject 41 — Space 1 (C5)

no description

### Subject 41 — Space 2 (C2)

Original text in French: “J’ai vu mon ex-conjoint devant moi. Il s’est présenté avec sa plus belle silhouette, il était tout noir et sont corps bougeait, le mouvement le faisait apparaitre et disparaitre. Je voyais des formes géométriques de plusieurs couleurs (rouge, jaune, bleu, mauve) qui s”emboitaient comme un vitrail en arrière plan en forme de d^ome. Vision d’une pieuvre fluorescente jaune.” Translation in English:”I saw my ex-partner in front of me. He appeared in his most beautiful form; he was entirely black, and his body was moving—this motion made him appear and disappear. I saw geometric shapes in various colors (red, yellow, blue, purple) interlocking like a stained glass window in the background, shaped like a dome. I had a vision of a fluorescent yellow octopus.”

### Subject 42 — Space 1 (C5)

“I heard a disembodied voice while in this space. It gave me the choice to “let go” and join them (there were others who were not pictured or heard) aka die, or I could go back to my body and return to earth to finish my journey.”

### Subject 42 — Space 2 (A7)

“No beings; the appearance of the space was pastel colored (light blues, light pinks, and light purples, and white – brighter white than is normal on earth); 3-d shaped, like I was floating upwards through a portal towards it.”

### Subject 42 — Space 3 (C2)

“The space felt like a horizon line; like I was standing on some kind of plane; There were other beings or people there, presumably lightbodies or souls/spirits. It was bright and light and the person or people around me communicated with me (non-verbally) and sent me so much love.”

### Subject 43 — Space 1 (C5)

no description

### Subject 43 — Space 2 (C5)

“1 God, appeared as a great light in the distance, the most important entity, corresponds to C5 in the form; 1 entity, square, followed me, a smaller light, was guiding me, my guardian angel ; Many entities, Angels, like the guardian angel following me, innumerable quantity, too many to count.”

### Subject 44 — Space 1 (C5)

“tall being glowing white robe long hair, could see his face but it was blurred. I could hear him talk in my mind, telepathically”

### Subject 45 — Space 1 (C5)

“objects were moving clockwise - life real from child to that moment in high resolution like a projector fast the size was normal ( like a scroll )”

### Subject 46 — Space 1 (C5)

”Outside space. Golden love light, rainbow (violent, pink, red, orange,yellow, white, teal blue green, purple more gold.) smudged it moving fast. counter clockwise. In the middle was a small black dot in center. That I would be visiting next. Cross, space 2. Plasma love light beings where made of plasma like lightening bolts.like a plasma ball. Thunder lines. They never stopped being part of me. It was like an electrical flower. It was really huge.”

### Subject 46 — Space 2 (C5)

“A black hole looks black from afar. The light was so bright it was hard to see all colors, they were still there. When I moved fast enough they all became very bright white. I was located in the middle. I was spinning in the center on the same axis. 360 degrees. Like a gyroscope I was turning the same way on the zero point of w, y, z.”

### Subject 47 — Space 1 (C5)

“It was clear perfectly color and I saw my dad very healthy and strong but he was very sick before he died”

### Subject 47 — Space 2 (B9)

“Self”

### Subject 47 — Space 3 (B10)

“I was happy enjoying the life in the second life and every one young and healthy and a lot of technology in their life and they moving to other places by time very fast like the light.”

### Subject 47 — Space 4 (B11)

“I felt the realty in the second life and it was clear vision and colorful”

### Subject 47 — Space 5 (B12)

“The same”

### Subject 48 — Space 1 (C5)

“I was in a black space that seemed to go on forever, yet felt enclosed, a sort of bubble. It felt infinite yet had a boundary. It was darkness yet seemed bright, no visible light source. I sensed 2 or 3 spirits to my left just outside the blackness - outside the membrane it seemed brighter - I could see them as if they were in a lighter space. I saw their head and shoulders, below their waits was hidden by the blackness. I felt they were there for me, cared about me, and I could trust them completely. I felt my body was horizontal facing down and supported everywhere, not just where I would be supported if I was laying on the ground. Many more beings began approaching from the left behind the original 2, and the membrane became thicker to protect me. It felt completely serene, calm, safe, suspended weightlessness.”

